# ApoE4 Promotes Thrombosis via Endothelial Cell ApoER2 and PP2A Activation

**DOI:** 10.64898/2026.08.17.745317

**Authors:** Yike Sun, Anastasia Sacharidou, Kenian Chen, Andrew Lemoff, Shiva Keshava, L. Vijaya Mohan Rao, Lin Xu, Chieko Mineo, Philip W. Shaul

## Abstract

**Background:** *APOE4*, the variant of apolipoprotein E carried by 25% of individuals, is a common genetic risk factor for cardiovascular disease (CVD). Although ApoE classically participates in lipid transport, *APOE4*-associated risk goes beyond impact on circulating lipids. Life-threatening CVD events including myocardial infarction and stroke are driven by atherogenesis and thrombosis. In mice ApoE4 increases atherosclerosis severity, but whether other major drivers of CVD events are influenced by ApoE4 is unknown.

**Methods:** GWAS data for venous thromboembolism (VTE) were analyzed. In humanized *APOE3* (hE3) and *APOE4* (hE4) mice, thrombosis was assessed by intravital microscopy (IVM) in the mesenteric microcirculation and by inferior vena cava (IVC) partial ligation. Actions of ApoE3 versus ApoE4 on endothelial cells (EC) and their underpinnings were studied in cultured human and mouse aortic EC, interrogating interactomes with immunoprecipitation-mass spectrometry and quantifying the secretion of Von Willebrand Factor (vWF), a critical initiator of thrombosis. Single cell transcriptomics datasets were queried do localize endothelial cell gene expression.

**Results:** GWAS showed that *APOE4* is associated with increased VTE risk, and whereas plasma lipids were similar, both microvascular and venous thrombosis were markedly increased in hE4 compared to hE3 mice. In cultured EC, whereas ApoE3 attenuated vWF secretion, it was enhanced by ApoE4, and both processes were mediated by ApoE receptor 2 (ApoER2). ApoE4, but not ApoE3, suppressed VEGF eNOS activation and NO production by causing the recruitment of the protein phosphatase 2A (PP2A) catalytic subunit to ApoER2 and the activation of PP2A. PP2A deletion prevented ApoE4-induced eNOS antagonism and vWF secretion by preserving Akt activation, and the NO donor spermine NONOate negated apoE4 stimulation of vWF secretion. PP2A activity was increased in hE4 aortas and IVC, and EC ApoER2 deletion or pharmacologic PP2A inhibition fully prevented exaggerated thrombosis in hE4 mice. In human great saphenous vein ApoER2 is primarily expressed in valvular endothelium.

**Conclusions:** *APOE4* is a risk allele for thrombosis, and ApoE4 is prothrombotic in microvasculature and veins in mice. Mechanistically, the ApoE4-EC ApoER2 tandem enhances vWF secretion by recruiting and activating PP2A and antagonizing eNOS, resulting in exaggerated thrombosis. In human veins ApoER2 is expressed in valvular endothelium, which is the most common site of initiation of venous thrombosis. Targeting these processes may afford protection from both primary thrombotic disorders like VTE and acute CVD events such as myocardial infarction and stroke in 25% of the population.

**CLINICAL PERSPECTIVE:** *What is New?:*

- Apolipoprotein E4 (ApoE4), a variant of ApoE carried by 25% of the population, increases cardiovascular disease risk independent of impact on circulating lipids, but the mechanism is unknown.
- This study reveals that through direct actions on endothelial cells, ApoE4 promotes thrombosis, which plays a critical role in acute cardiovascular events like myocardial infarction and stroke.
- Our findings demonstrate that ApoE4 uniquely causes ApoE receptor 2 (ApoER2) in endothelial cells to recruit and activate the enzyme protein phosphatase 2A (PP2A), leading to exaggerated secretion of Von Willebrand Factor (vWF), which is a major initiator of thrombosis.

*What Are the Clinical Implications?:*

- We demonstrate that ApoE4 promotes both microvascular and venous thrombosis, and that ApoER2 and PP2A are expressed in human leg vein endothelium that are instrumental in the initiation of venous thromboembolism, which has a 30-day case fatality rate of up to 30%.
- Until now there have been no interventions available to break the link between ApoE4 and life-threatening vascular disorders, and we demonstrate that pharmacologic inhibition of PP2A fully prevents the exaggerated thrombosis caused by ApoE4.
- Targeting the mechanisms initiated by ApoE4 and ApoER2 in the endothelium may afford protection from both primary thrombotic orders like VTE and acute cardiovascular events such as myocardial infarction and stroke in 25% of the population.

## INTRODUCTION

Apolipoprotein E (ApoE) is a 34 kDa protein composed of 299 amino acids that is primarily produced in the liver^1^. ApoE classically participates in lipid transport by binding to members of the LDL receptor family^2^. The three common isoforms of ApoE in humans (ApoE2, ApoE3 and ApoE4) are encoded by a single gene on chromosome 19, and they are distinguished by single point variations at positions 112 and 158; ApoE2 has Cys112 and Cys158, ApoE3 has Cys112 and Arg 158, and ApoE4 has Arg 112 and Arg158. In ApoE4 the presence of Arg112 leads to C-terminal-to-N-terminal domain folding that underlies its altered functions compared with ApoE2 and ApoE3^3–5^. The allele frequencies are 5-10% for *APOE2*, 65-70% for *APOE3*, and 15-20% for *APOE4*^6^. *APOE4* is a genetic risk factor for cardiovascular disease (CVD). Compared to individuals homozygous for *APOE3*, those harboring the *APOE4* allele have an increased risk of coronary artery disease (CAD) and poorer survival following myocardial infarction^7–10^. Although the impact of ApoE4 may partially relate to its effect on plasma cholesterol and LDL abundance^5^, multiple studies indicate that ApoE4-associated vascular disease risk goes well beyond influence on lipoprotein status^8,11–13^. This includes evidence that despite similar lipid status on statins, atherosclerosis-related vascular complications occur more frequently in individuals with *APOE4*^14^. Along with its impact on coronary and peripheral artery disease risk, *APOE4* increases the likelihood of cerebrovascular disease^15,16^.

The two primary pathogenetic processes that lead to the major life-threatening clinical events of CVD, namely myocardial infarction and stroke, are atherogenesis and thrombosis^17,18^. In the backdrop of atherosclerosis development, repeated plaque microruptures cause subclinical thrombosis that promotes plaque growth and vulnerability^19^. Thrombosis also plays a critical role in the dire clinical presentations because acute thrombus formation following the rupture or erosion of an atherosclerotic plaque is a major cause of acute coronary syndrome, myocardial infarction, transient ischemic attack and ischemic stroke^20,21^. In addition to the contribution of thrombosis to atherogenesis and related acute clinical CVD events, there are distinct thrombotic disorders like venous thromboembolism (VTE). VTE includes pulmonary embolism (PE) and deep vein thrombosis (DVT), and has a 30-day case fatality rate of 11-30% in the general population^22–27^. Whereas both human and mouse studies have indicated that atherosclerosis severity is increased in the setting of ApoE4 ^28–32^, whether ApoE4 promotes thrombosis is unknown.

Despite ApoE4 being designated as a unique isoform of ApoE over 40 years ago^33^, how ApoE4 increases CVD risk remains poorly understood, and there are no interventions available to break the link between *APOE4* and life-threatening vascular disorders. To fill these important knowledge and therapeutic gaps, in the current work experiments were designed to determine if and how ApoE4 promotes thrombosis. They revealed that *APOE4* is associated with greater risk of VTE in a multipopulation GWAS in which eight cohorts have been combined, and employing two entirely independent models, thrombosis was markedly increased in humanized ApoE4 mice compared to ApoE3 mice. Efforts then turned to the interrogation of the underlying mechanisms, revealing that the underpinnings are endothelial in nature, involving the LDL receptor family member apolipoprotein E receptor 2 (ApoER2) and recruitment and activation of the serine/threonine protein phosphatase 2A (PP2A) leading to eNOS antagonism and the promotion of endothelial cell Von Willebrand Factor (vWF) secretion. These findings reveal an entirely new mode of action of ApoE4 that may represent a set of targets to leverage to combat both primary thrombotic disorders like VTE and acute CVD events such as myocardial infarction and stroke in 25% of the population. That possibility is demonstrated by the discovery that administration of the PP2A inhibitor Endothall fully protects ApoE4 mice from exaggerated thrombosis.

## METHODS

Additional detailed descriptions of the methods are available in the Supplemental Materials.

### GWAS Analyses

The association between *APOE4* and venous thromboembolism (VTE) was evaluated in 6355 VTE patients and 161,416 non-VTE controls from eight cohorts^34^. To provide an assessment of the link between *APOE4* and other cardiovascular disorders, we also performed association analysis for additional conditions. These phenotype-specific associations were summarized descriptively in a sun plot, recognizing that it was necessary to derive the estimates from multiple GWAS cohorts rather than from a single unified study population.

### Animal Models

In vivo studies were performed in mice in which the endogenous *Apoe* gene has been replaced with the human *APOE3* or *APOE4* gene at the same locus^31,35^. The mice are referred to as hE3 and hE4, respectively. To evaluate the role of the LDL receptor family member ApoER2, which is encoded by *LRP8*, in the endothelium, ApoER2 floxed mice that we previously generated (ApoER2^fl/fl^)^36^ were crossed with VE-cadherin Cre mice (VECad-Cre)^37^, and the resulting control mice (ApoER2^fl/fl^) and those selectively deficient in ApoER2 in endothelium (ApoER2^ΔEC^) were placed on hE3 or hE4 background. In additional experiments hE3 and hE4 mice were treated with the PP2A inhibitor Endothall^36^.

### Thrombosis Models

To evaluate microvascular thrombosis intravital microscopy was employed in the mesenteric microcirculation as previously described^36^. To study venous thrombosis, a partial flow restriction (stenosis) model in the inferior vena cava (IVC) was used^38^.

### Cell Culture Experiments

Studies were performed in primary human aortic endothelial cells (HAEC) and mouse aortic endothelial cells (MAEC). Gene knockdown was accomplished by shRNA using lentiviral constructs. Recognizing that von Willibrand Factor (vWF) release from endothelial cells through the exocytosis of Weibel-Palade bodies (WPB) is a major initiating step in thrombosis^39,40^, the effects of ApoE3 and ApoE4 on vWF secretion were evaluated in HAEC and in MAEC. Endothelial cell NO synthase (eNOS) activity was evaluated by the measurement of total nitrate and nitrite in the media, and the assessment of eNOS-Ser1177 phosphorylation. Akt activation was also evaluated by immunoblotting for phosphorylated Akt-Ser473. Protein co-immunoprecipitation was done by previously-described methods^41^, and immunoblotting was additionally performed to confirm effective gene knockdown.

### Liquid Chromatography/ Tandem Mass Spectrometry (LC/MS-MS)

The ApoER2 interactome in endothelial cells and its dynamic changes in response to ApoE3 versus ApoE4 were evaluated by liquid chromatography/tandem mass spectrometry (LC/MS-MS) using methods modified from prior approaches^41^.

### PP2A Activity

PP2A phosphatase activity in cultured endothelial cells or mouse blood vessels was evaluated using PP2A immunoprecipitation, incubation with a phosphorylated peptide substrate, and the quantification of released free phosphate^36^.

### Vascular Cell Transcriptome Analyses

To interrogate the vascular cell distribution of expression of LRP8, ApoE and PP2A subunit genes, four publicly available single cell RNA-seq datasets were queried^42–45^.

### Statistical Analysis

Following normality testing by a Shapiro—Wilk test, for normally-distributed datasets comparisons between two groups were performed by two-sided, unpaired Welch’s or Student’s t tests, and differences between more than two groups were evaluated by one-way analysis of variance (ANOVA) with Tukey’s post-hoc testing. In the instances in which normality testing failed, non-parametric analyses were performed between two groups using Mann—Whitney U test, and between more than two groups by non-parametric ANOVA with Kruskal—Wallis and Dunn’s post-hoc testing. Chi-square testing was employed to compare observed frequencies in categorical data. Values are reported as mean ± SEM. Significance was accepted at the 0.05 level of probability.

## RESULTS

### ApoE4 is a risk allele for thrombosis

To determine whether *APOE4* is associated with thrombosis risk, we interrogated GWAS data from 6,355 VTE cases and 161,416 non-VTE controls across eight cohorts^34^. The association between *APOE4* and other cardiovascular phenotypes was also assessed (Table S1). Because it was necessary to derive estimates from distinct GWAS datasets, direct cross-trait effect-size comparison is not feasible and the findings are presented as a descriptive cross-phenotype summary in the sun plot in Figure 1. The analyses revealed that along with being associated with other cardiovascular phenotypes, *APOE4* is a risk allele for VTE (P = 0.0042). This finding across multiple cohorts provides genetic evidence that compared with the more common *APOE3* variant, *APOE4* is associated with increased risk of VTE.

**Figure 1.**
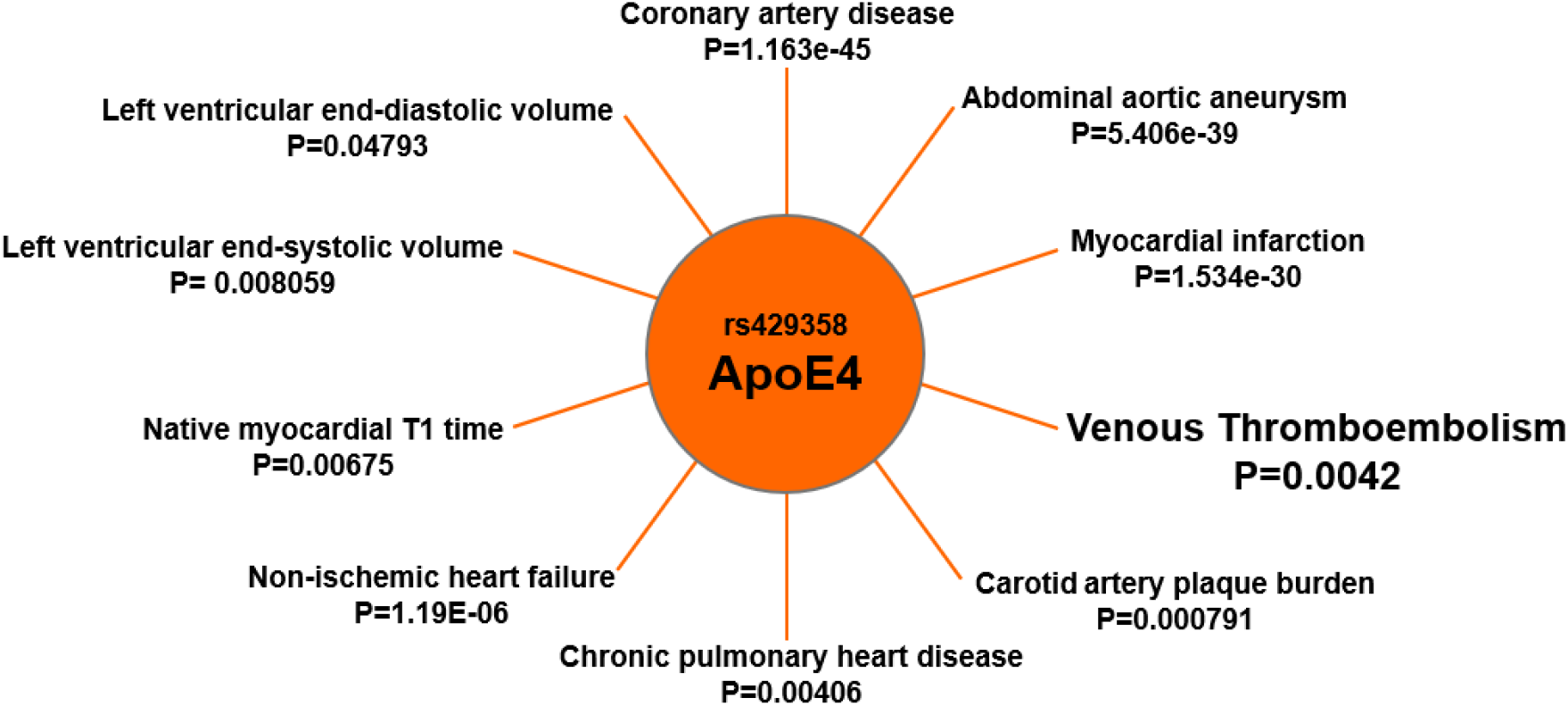
ApoE4 is associated with thrombosis risk and other cardiovascular phenotypes. The sun plot summarizes associations between rs429358/*APOE4* and VTE and other cardiovascular phenotypes, including coronary artery disease, abdominal aortic aneurysm, myocardial infarction, carotid artery plaque burden, chronic pulmonary heart disease, non-ischemic heart failure, left ventricular end-diastolic and end-systolic volumes, and native myocardial T1 time. P values are shown for each phenotype. Because it is necessary to derive estimates from distinct GWAS datasets, direct cross-trait effect-size comparison is not feasible and the findings are presented as a descriptive cross-phenotype summary. See Table S1.

### ApoE4 promotes thrombosis

To directly determine if ApoE4 enhances thrombosis, we performed intravital microscopy and quantified thrombus formation in the mesenteric microcirculation of humanized hE3 and hE4 mice (Figure 2A-C, and Videos S1 and S2). The size of the largest thrombus formed, within 7 min for arterioles and within 3 min for venules, and the time to full occlusion following the initiation with ferric chloride application were quantified. In hE4 mice the maximum sized thrombus was increased by 9.9-fold in arterioles and by 39-fold in venules (Figure 2D,E). The time to full occlusion in arterioles was decreased by 48%, and in venules it was decreased by 65% (Figure 2F,G). Thus, ApoE4 promotes thrombosis in the microcirculation, in both arterioles and venules.

**Figure 2.**
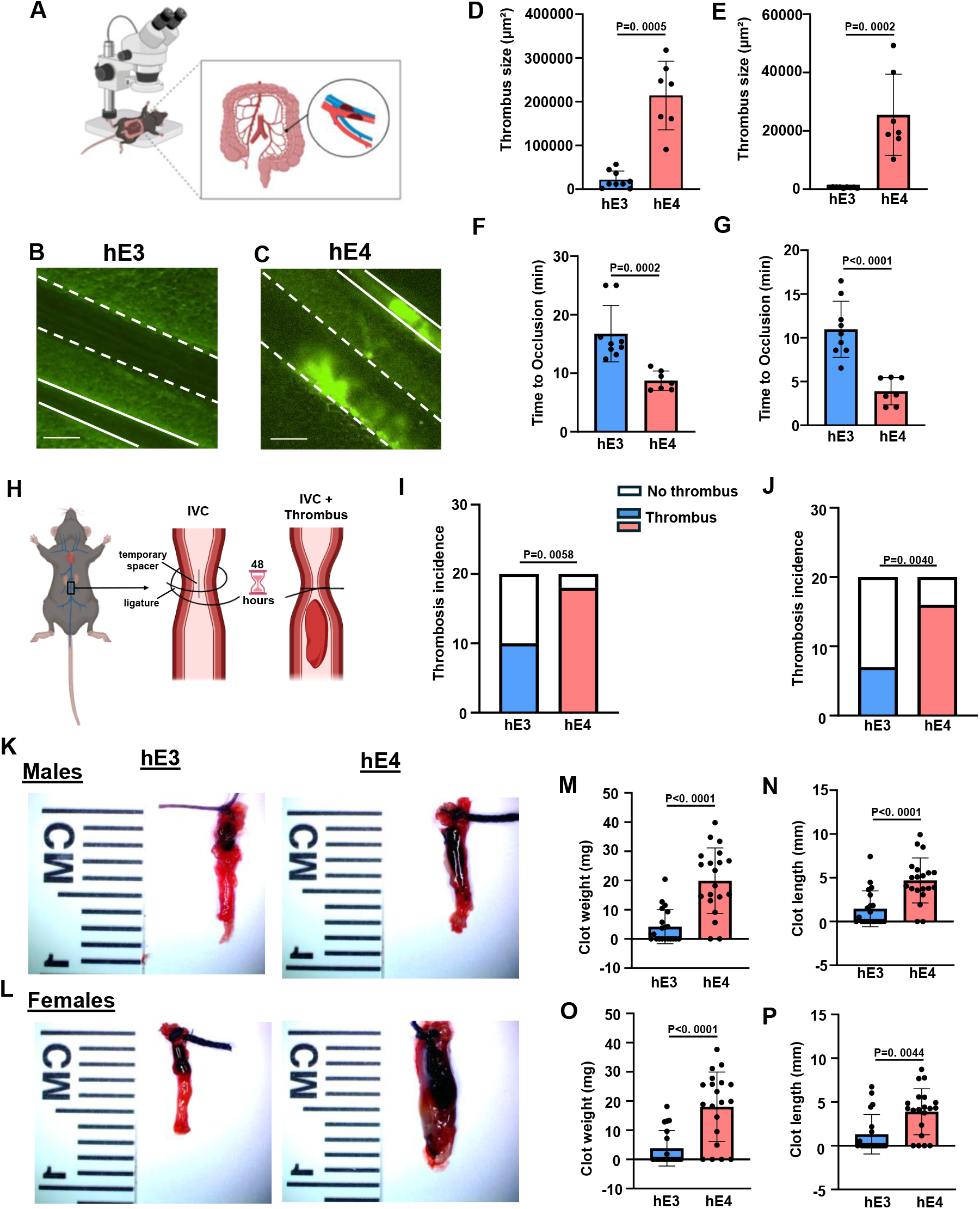
ApoE4 promotes microvascular and venous thrombosis. **A-G**, Thrombus formation in mesenteric arterioles and venules was compared in hE3 versus hE4 mice by intravital microscopy (**A**). Platelets were fluorescence-labeled and ferric chloride was applied to the mesentery to initiate thrombosis. **B,C,** Representative still images, with arteriole walls highlighted by dashed lines and venule walls highlighted by solid lines. Scale bar=50um. **D, E**, The size of the largest thrombus observed following ferric chloride application was quantified in arterioles (**D,** within 7 min, n=9 and 7 per group) and venules (**E**, within 3 min, n=9 and 7 per group). **F, G**, The time to full occlusion of the arterioles (**F**) and venules (**G**) was quantified (n= 9 and 7 per group). **H-P**, A partial flow restriction (stenosis) model of thrombosis was tested in the inferior vena cava (IVC). A ligature is placed around the IVC inferior to the renal veins, using a temporary spacer that is removed to maintain a lumen area of 10% (**H**). The IVC is harvested 48h later. **I, J,** The incidence of thrombus formation was evaluated in male (**I)** and female mice (**J). K, L,** Representative images of the thrombus in the dissected inferior vena cava in male (**K**) and female mice (**L**). **M, N**, Clot weight (**M**) and length (**N**) in male mice (n=20 per group). **O, P**, Clot weight (**O**) and length (**P**) in female mice (n=20 per group). Data normality was evaluated by Shapiro-Wilk tests. Incidence of thrombosis in the IVC model was compared by Chi-square test (**I** and **J**). Values are mean±SEM. For normally distributed datasets the comparisons between two groups were evaluated by two-sided, unpaired Welch’s t test (**D** and **G**), In the instances in which normality testing failed, non-parametric analyses were performed between two groups using Mann—Whitney U test (**E**, **F**, **M-P**).

To provide a model of venous thrombosis, a partial flow restriction (stenosis) model was employed in the mouse inferior vena cava (IVC) (Figure 2H). A ligature is placed around the IVC inferior to the renal veins, using a temporary spacer that is removed to maintain a lumen area of 10%. The IVC is harvested 48h later. The incidence of thrombosis was increased to 90% from 50% in hE4 compared to hE3 male mice (Figure 2I), and it was increased to 80% from 35% in hE4 compared to hE3 female mice (Figure 2J). In both males and females plasma total cholesterol, triglycerides, LDL cholesterol and HDL cholesterol concentrations were similar in hE3 and hE4 mice (Figure S1). Representative images of IVC and thrombi therein are provided in Figure 2K and 2L. The clot weight and length were increased by 477% and 321%, respectfully, in male hE4 mice (Figure 2M,N), and similarly they were increased by 474% and 296%, respectfully, in female hE4 mice (Figure 2O,P). As such, in the absence of differences in circulating lipids, ApoE4 causes robust enhancement of both microvascular and venous thrombosis.

### The ApoE4-ApoER2 tandem stimulates endothelial cell vWF secretion

Recognizing that von Willibrand Factor (vWF) release from endothelial cells through the exocytosis of Weibel-Palade bodies (WPB) is a major initiating step in thrombosis^46^, the effects of ApoE3 and ApoE4 on vWF secretion were evaluated in HAEC (Figure 3A). Predictably, thrombin increased vWF secretion, and VEGF attenuated the secretion prompted by thrombin (Figure 3B). Whereas ApoE3 also blunted vWF secretion in response to thrombin, in contrast ApoE4 caused an increase in secretion (Figure 3C). The roles of various ApoE receptors expressed in endothelial cells were then evaluated. In the setting of LDLR knockdown the attenuation of thrombin-stimulated vWF secretion by ApoE3 was preserved, and the enhancement in secretion in response to ApoE4 was also maintained (Figure 3D, Figure S2A). That was also the case with VLDLR knockdown (Figure 3E, Figure S2B) or loss of LRP-1 (Figure 3F, Figure S2C). However, in the setting of ApoER2 knockdown the decrease in thrombin-stimulated vWF secretion by ApoE3 was absent, and ApoE4 also had no effect (Figure 3G,H, Figure S2D,E). These findings reveal that ApoE3 and ApoE4 have opposing effects on endothelial cell vWF release, being anti-thrombotic and pro-thrombotic in nature, respectively, and that endothelial cell ApoER2 is required for both processes.

**Figure 3.**
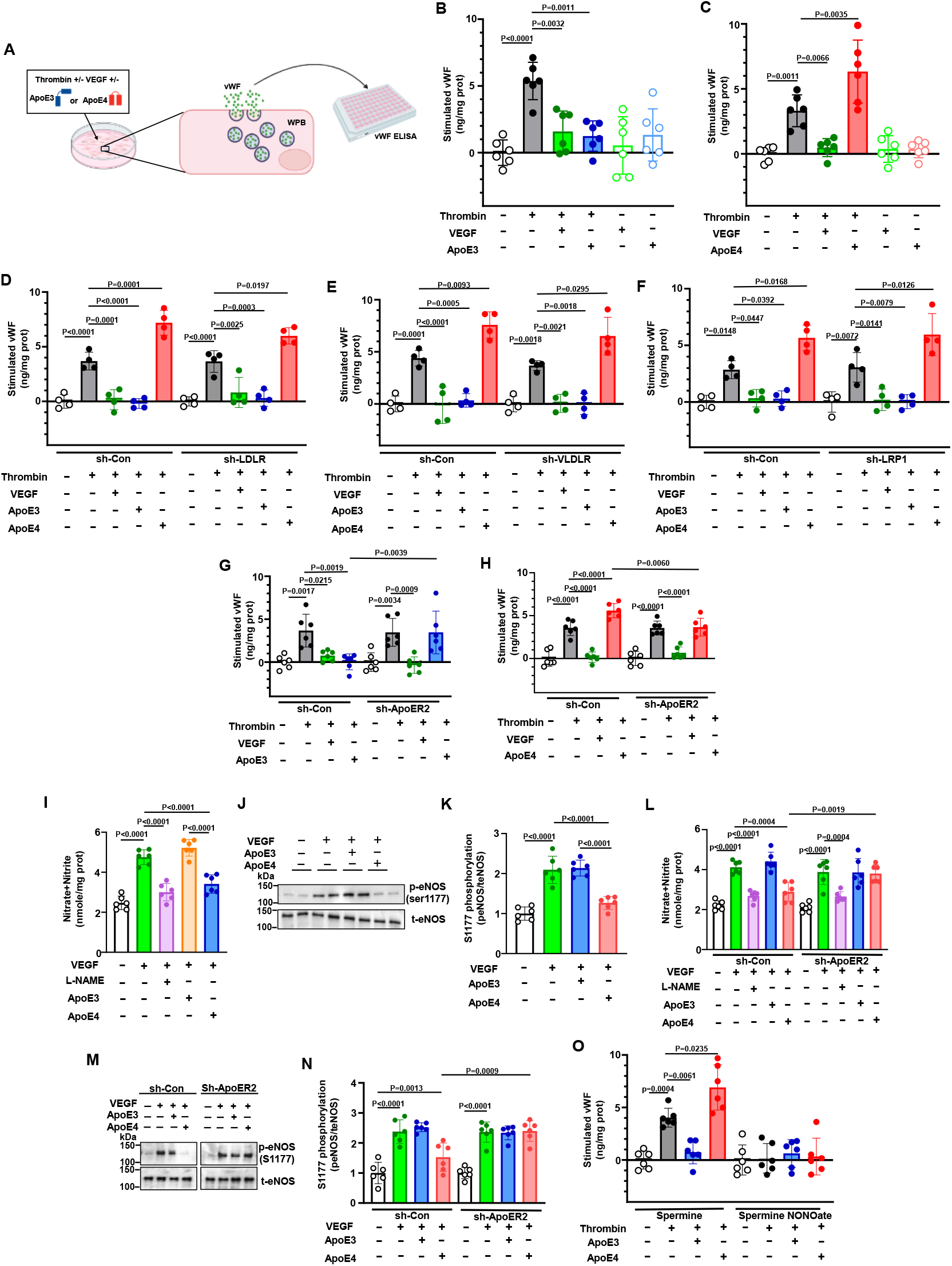
ApoE4 activates PP2A via ApoER2 to antagonize eNOS and enhance vWF secretion. **A,** vWF secretion from HAEC was measured in the absence or presence of thrombin (3U/ml), using VEGF (100ng/ml) treatment as a control for attenuation of secretion. Values shown are relative to basal abundance in the absence of treatment. **B**, The effect of ApoE3 (15ug/ml) was evaluated, n=6/group. **C**, The effect of ApoE4 (15ug/ml) was evaluated, n=6/group. **D-H**, ApoE3 and ApoE4 modulation of vWF secretion was tested in the setting of control shRNA or shRNA knockdown of the apoE receptors LDLR (**D**, n=4/group), VLDLR (**E**, n=4/group), LRP1 (**F**, n=4/group), or ApoER2 (**G,H**, n=6/group). **I,** NO production by HAEC was determined by the measurement of total nitrate and nitrite in the media, following treatment with vehicle or VEGF (100ng/ml) in the absence or presence of L-NAME (2mM), ApoE3 or ApoE4 (15ug/ml). N=6/group. **J K,** Phosphorylated eNOS Ser-1177 and total eNOS were detected by immunoblotting following HAEC treatment with vehicle or VEGF in the absence or presence of ApoE3 or ApoE4. Representative immunoblots with two samples per treatment are in **J**, and quantification is in **K** (n=6/group). **L**, NO production with vehicle or VEGF (100ng/ml) in the absence or presence of L-NAME, ApoE3 or ApoE4 was quantified after control siRNA or siRNA knockdown of ApoER2. N=6/group. **M, N**, Activating eNOS Ser-1177 phosphorylation was evaluated in HAEC treated with vehicle or VEGF plus/minus ApoE3 or ApoE4 after control shRNA or shRNA knockdown of ApoER2. Representative immunoblots with one sample per treatment are in **M**, and quantification is in N (n=6/group). **O**, vWF secretion was measured in the absence or presence of thrombin plus/minus ApoE3 or ApoE4, in cells concurrently treated with the NO donor spermine NONOate or the parent compound spermine (1uM). N=6/group. Values are mean±SEM. Datasets were normally distributed, and groups were compared by analysis of variance (ANOVA) with Tukey’s post-hoc testing.

### ApoE4 activates PP2A via ApoER2 to antagonize eNOS and enhance vWF secretion

One of the major modes of regulation of vWF secretion through the exocytosis of Weibel-Palade bodies (WPB) involves the modulation of the activity of N-ethylmaleimide-sensitive factor (NSF) by nitric oxide (NO). The nitrosylation of key cysteines attenuates NSF disassembly of soluble NSF attachment protein receptor (SNARE) complexes, thereby decreasing the exocytosis of WPB^47^. We therefore evaluated the effects of ApoE3 and ApoE4 on VEGF activation of eNOS in HAEC by quantifying NO production by measurement of total nitrate and nitrite release and the activating phosphorylation of eNOS-Ser1177 (Figure S3). Whereas ApoE3 had no effect, ApoE4 fully prevented NO production and the underlying required eNOS-Ser1177 phosphorylation in response to VEGF (Figure 3I-K). The antagonist action of ApoE4 on NO production and eNOS activation was entirely dependent on ApoER2 (Figure 3L-N). The exogenous NO donor spermine NONOate prevented the enhancement of vWF secretion by ApoE4 (Figure 3O), indicating that the antagonism of eNOS by ApoE4 underlies its stimulation of vWF release.

To determine how the ApoE4-ApoER2 tandem inhibits eNOS leading to effects on vWF secretion, we leveraged the differential effects of ApoE3 versus ApoE4 and interrogated the ApoER2 interactome in endothelial cells incubated with ApoE3 and ApoE4. HAEC were treated with vehicle, ApoE3 or ApoE4 for 15 min, ApoER2 was immunoprecipitated, and liquid chromatography/tandem mass spectrometry (LC/MS-MS) was performed (Figure 4A). It was found that 675 proteins were more abundantly associated with ApoER2 in response to ApoE3 treatment and 610 proteins were more prominent in the ApoER2 interactome in response to ApoE4 (Table S2). The proteins modulated differentially by ApoE4 compared to ApoE3 included subunits of protein phosphatase 2A (PP2A). It is a serine/threonine protein phosphatase comprised of a trimeric protein complex in which a core dimer formed between the scaffolding A subunit (either PPP2R1A or PPP2R1B) and the catalytic C subunit (either PPP2CA or PPP2CB) is associated with one of the many B subunits (including PPP2R2A) that facilitate and direct the interaction of the trimer with substrate proteins ^48^. PPP2R1A, PPP2R2A and PPP2CA, representing A, B and C subunits of the phosphatase respectively, were all recruited to ApoER2 in response to ApoE4 compared to in response to ApoE3 (Table S2). The ApoE4-induced recruitment of PP2A subunits to ApoER2 has possible mechanistic significance because PP2A mediates the dephosphorylation of eNOS and Akt, which is the kinase responsible for eNOS-Ser1177 phosphorylation in response to many agonists^49^.

**Figure 4.**
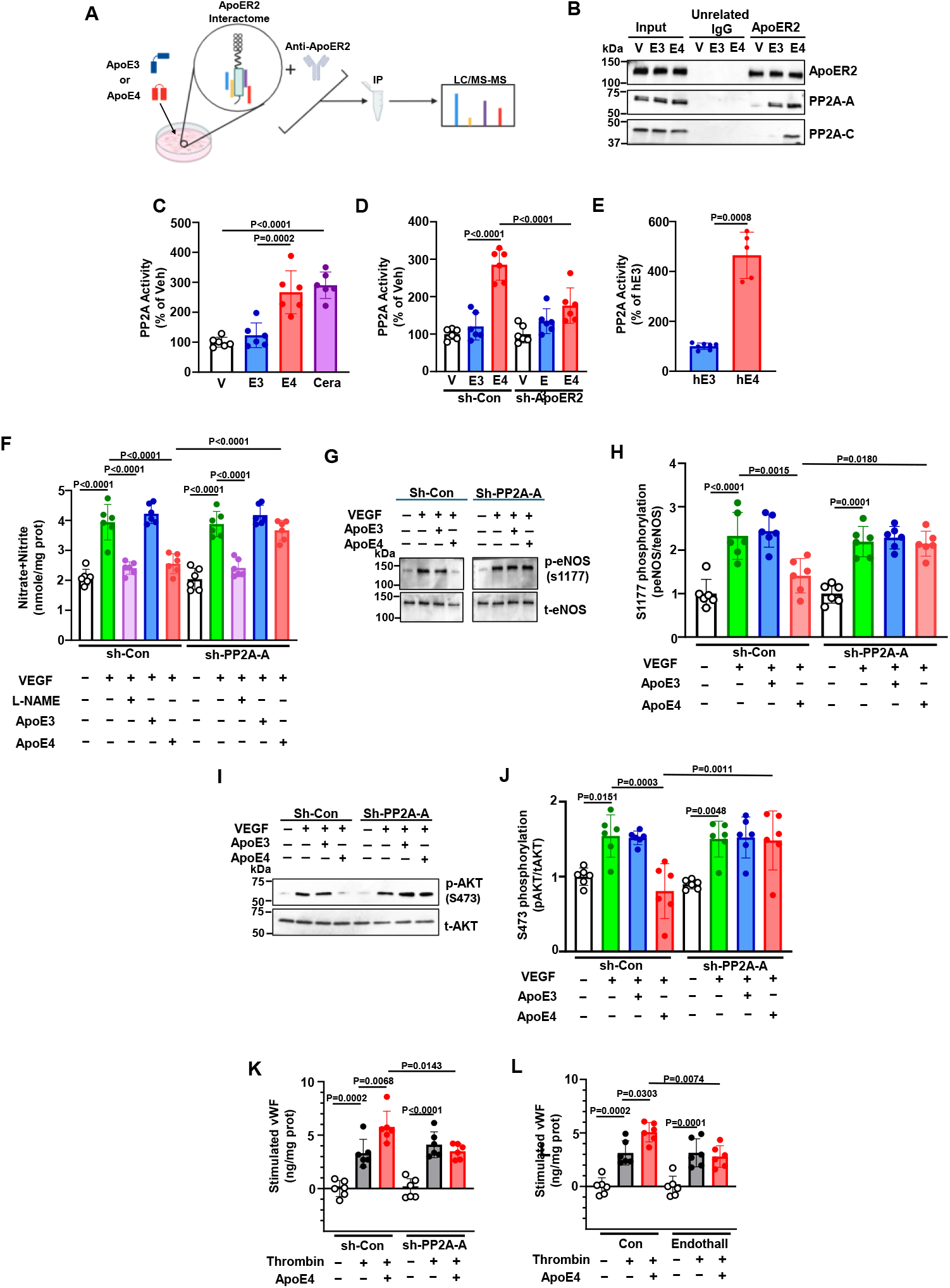
ApoE4 causes PP2A recruitment to ApoER2 and PP2A activation, leading to Akt and eNOS dephosphorylation and decreased NO production, which results in exaggerated vWF secretion. **A,** The ApoER2 interactome was interrogated in HAEC treated with ApoE3 or ApoE4 by immunoprecipitation (IP) and liquid chromatography and tandem mass spectrometry (LC/MS-MS). **B,** HAEC were treated with vehicle (V), ApoE3 or ApoE4, co-immunoprecipitation was performed with mock IgG or anti-ApoER2 antibody, and immunoblotting was performed to detect ApoER2, PP2A-A or PP2A-C. **C,** PP2A enzymatic activity was measured in HAEC treated with vehicle, ApoE3 or ApoE4 (15ug/ml), or ceramide positive control (Cera, 30 uM). N=6. **D**, A parallel experiment was performed after control shRNA or shRNA knockdown of ApoER2. N=6. **E**, PP2A enzymatic activity was measured in aortas obtained from hE3 and hE4 mice. N=7 and 5, respectively. **F**, NO production with vehicle or VEGF (100ng/ml) in the absence or presence of L-NAME (2mM), ApoE3 or ApoE4 was quantified after control shRNA or shRNA knockdown of PP2A-A. N=6/group. **G,H**, Activating eNOS Ser-1177 phosphorylation was evaluated in HAEC treated with vehicle or VEGF plus/minus ApoE3 or ApoE4 after control shRNA or shRNA knockdown of PP2A-A. Representative immunoblots with one sample per treatment are in **G**, and quantification is in **H** (n=6/group). **I,J**, Activating Akt Ser-473 phosphorylation was evaluated in HAEC treated with vehicle or VEGF plus/minus ApoE3 or ApoE4 after control shRNA or shRNA knockdown of PP2A-A. Representative immunoblots with one sample per treatment are in **I**, and quantification is in **J** (n=6/group). **K, L**, vWF secretion was measured in the absence or presence of thrombin plus/minus ApoE4, after control shRNA or shRNA knockdown of PP2A-A (**K**), or in cells concurrently treated with the PP2A inhibitor endothall or the control compound 1,4-dimethyl endothall (1uM) (**L**). N=6/group. Values are mean±SEM. Datasets were normally distributed, and groups were compared by either two-sided, unpaired Welch’s t test (**E**) or analysis of variance (ANOVA) with Tukey’s post-hoc testing (**C, D, F, H, J, K-L**).

Prompted by the findings about the ApoER2 interactome made by LC/MS-MS, co-immunoprecipitation was performed to assess PP2A recruitment to ApoER2 in response to ApoE3 and ApoE4 (Figure 4B). With both forms of ApoE there was recruitment of PP2A-A, but it was only in response to ApoE4 that PP2A-C was additionally recruited. Whether PP2A enzymatic activity is affected by ApoE3 or ApoE4 was evaluated using Malachite green. Whereas HAEC treatment with ApoE3 yielded no change in PP2A activity, ApoE4 caused a 2.7-fold increase in PP2A activity, and the activation was ApoER2-dependent (Figure 4C,D). To determine if PP2A activation occurs in vivo in response to ApoER4, the activity in aorta was quantified, and it was more than 4-fold greater in aortae from hE4 mice compared to hE3 mice (Figure 4E).

The role of PP2A in the effects of ApoE4 on eNOS was then assessed, and both the attenuation of NO production and the antagonism of activating eNOS-1177 phosphorylation by ApoE4 were prevented by PP2A-A excision (Figure 4F-H, respectively, and Figure S2F). Since Akt mediates the activating phosphorylation of eNOS^49^, its activation in response to VEGF was also evaluated by immunoblotting. ApoE4 blunted the activating phosphorylation of Akt, and the negative impact was prevented by knockdown of PP2A-A (Figure 4I,J). In addition, the exaggeration of vWF secretion with thrombin by ApoE4 was negated by either PP2A-A silencing or the PP2A antagonist endothall (Figure 4K,L). Collectively these observations indicate that the ApoE4 and ApoER2 partnership promotes endothelial cell vWF secretion by recruiting and activating PP2A, leading to the antagonism of Akt and eNOS and the loss of NO blunting of WPB exocytosis.

### ApoER2 and PP2A are expressed in human arterial, venous and valvular endothelium, and upregulated with thrombosis

Expanding on the discovery of a partnership between ApoER2 and PP2A in cultured endothelial cells, we interrogated a single cell transcriptomics dataset that interrogated vascular cells in 19 human tissues (Figure 5A-C)^42^. The UMAP plot in Figure 5A depicts the endothelial cell clusters in various human organs, using the group identities established by Barnett and colleagues^42^. The distribution of expression of genes relevant to ApoE4-ApoER2-induced thrombosis is shown in dot blots (Figure 5B,C). LRP8, which encodes ApoER2, is most highly expressed in the human blood-brain barrier endothelium, with addition expression in the brain venous endothelium, aorta and coronary artery endothelium, other arterial endothelium (art_ec_1), venous endothelium categorized in the original work to be expressed in most organs (ven_ec_1), and in venous endothelium of skeletal muscle, adipose tissue and lymph nodes (ven_ec_2). Regarding PP2A, PPP2R1A is the primary A subunit in human endothelial cells in situ. The C subunit genes PPP2CA and PPP2CB are abundant in blood-brain barrier, brain venous, other venous, and arterial endothelial cells, and lower expression is observed in capillary endothelium. The B subunit designated PPP2R2A is abundant in brain arterial and venous endothelium and blood-brain-barrier endothelium, and in other venous endothelial cells. The B subunit PPP2R5C, A and E are also relatively prevalent in endothelial cells compared to other B subunit genes.

**Figure 5.**
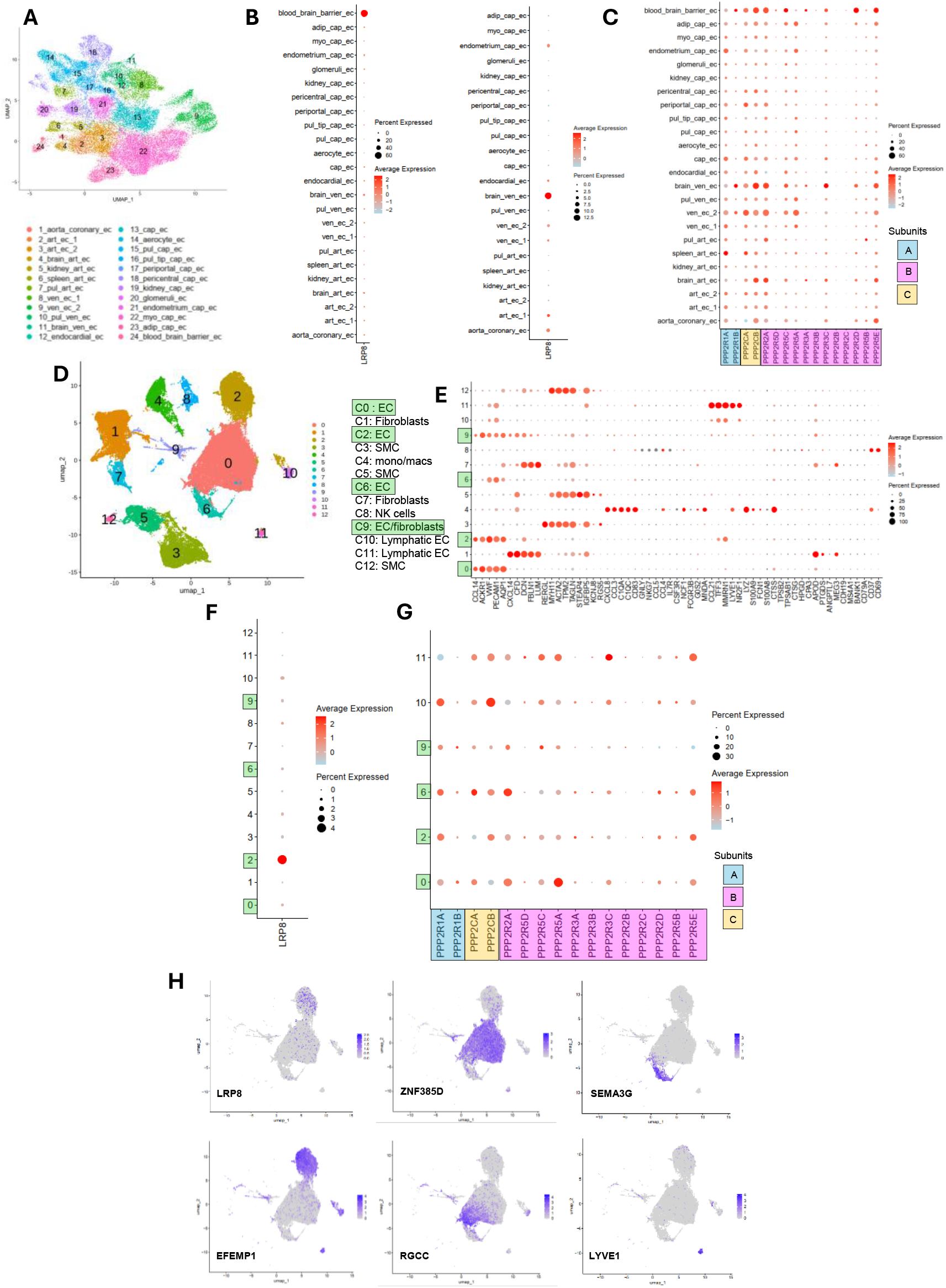
ApoER2 (LRP8) and PP2A are expressed in human arterial, venous and valvular endothelium. A, UMAP plot of endothelial cell (EC) populations in human organs. **B,C**, Dot plots for expression of LRP8 (**B**) and PP2A subunit subtypes (**C**) in the EC populations. **D, E,** UMAP plot of cell type clusters in the human great saphenous vein (**D**) and marker gene expression summary (**E**). **F**, Dot plot for expression of LRP8 in the various venous cells. **G**, Dot plot for expression of PP2A subunits in EC populations. **H**, UMAP plots for EC subtypes displaying the distribution of expression of LRP8 and of ZNF385D, SEMA3G, EFEMP1, RGCC and LYVE1 to identify venous, arterial, valvular, capillary and lymphatic EC, respectfully.

Noting that thrombosis commonly occurs in lower extremity veins, ApoER2 and PP2A expression in the great saphenous vein was interrogated using a publicly available dataset^44^. Marker gene analysis identifies clusters C0, C2, C6 and C9 as blood endothelial cells (Figure D,E). LRP8 is expressed predominantly in the endothelial cells (Figure 5F), and the endothelium express primarily PPP2R1A as the A subunit, and PPP2R2A, PPP2R5A and PPP2R5E are the principal B subunits (Figure 5G). Using markers for endothelial cell subtypes, such as ZNF385D, SEMA3G, EFEMP1, RGCC and LYVE1 to identify venous, arterial, valvular, capillary and lymphatic endothelial cells, respectively (Figure 5H), UMAP plots for the endothelial cell populations reveal that LRP8 is primarily expressed in valvular endothelium and also found in venous endothelium.

The possible impact of thrombosis on venous LRP8 and PP2A expression was evaluated in data from a prior study in the mouse IVC^45^. Marker gene expression designates cluster C5 as endothelial cells (Figure S4A,B). Lrp8 is most abundant in the endothelium (Figure S4C), in the endothelium Ppp2r1a is the primary A subunit, there are a variety of specific B subunits with Pppr5a being the most abundant, and both Ppp2ca and Ppp2cb are C subunits. Compared to sham instrumented IVC, with thrombosis endothelial cell Lrp8 increases by 61%, PP2A B subunits Ppp2r3c and Ppp2r2a increase by 2.2-fold and 55%, respectively, and PP2A C subunit Ppp2cb increases by 2.0-fold (Figure S4E). Collectively these findings in human and mouse vasculature reveal that ApoER2 and PP2A are expressed in human arterial, venous and valvular endothelium, and that the endothelial cell mechanisms driving ApoE4 promotion of thrombosis may actually be amplified in the setting of thrombosis.

### Endothelial cell ApoER2 and PP2A mediate ApoE4 promotion of thrombosis in vivo

Having implicated ApoER2 in the stimulation of vWF secretion by ApoE4 in HAEC in culture, to determine the role of the endothelial receptor in thrombosis in vivo we first assessed ApoE4 impact on vWF release from cultured mouse aortic endothelial cells (MAEC). Paralleling the observations in HAEC, in MAEC ApoE3 blunted vWF secretion and it was enhanced by ApoE4 (Figure 6A), and both responses were dependent on ApoER2. The partial flow restriction (stenosis) model of thrombosis in the IVC was then studied in male hE3 and hE4 mice with normal endothelial cell ApoER2 expression (ApoER2^fl/fl^) or deficient in endothelial ApoER2 (ApoER2^ΔEC^). In mice expressing endothelial cell ApoER2 the incidence of thrombosis was increased from 60% to 85% in hE4 compared to hE3 (p=0.1552), but there was negligible difference in incidence in mice lacking endothelial cell ApoER2 (Figure 6B). Representative images of IVC containing clots are in Figure 6C. The clot weight and length were increased by 339% and 311%, respectfully, in hE4 versus hE3 mice harboring normal endothelial cell ApoER2 (Figure 6D,E). In contrast, clot weight and length were similar in ApoER2^ΔEC^ mice on hE3 versus hE4 background. Thus, endothelial cell ApoER2 mediates ApoE4 promotion of thrombosis in vivo.

**Figure 6.**
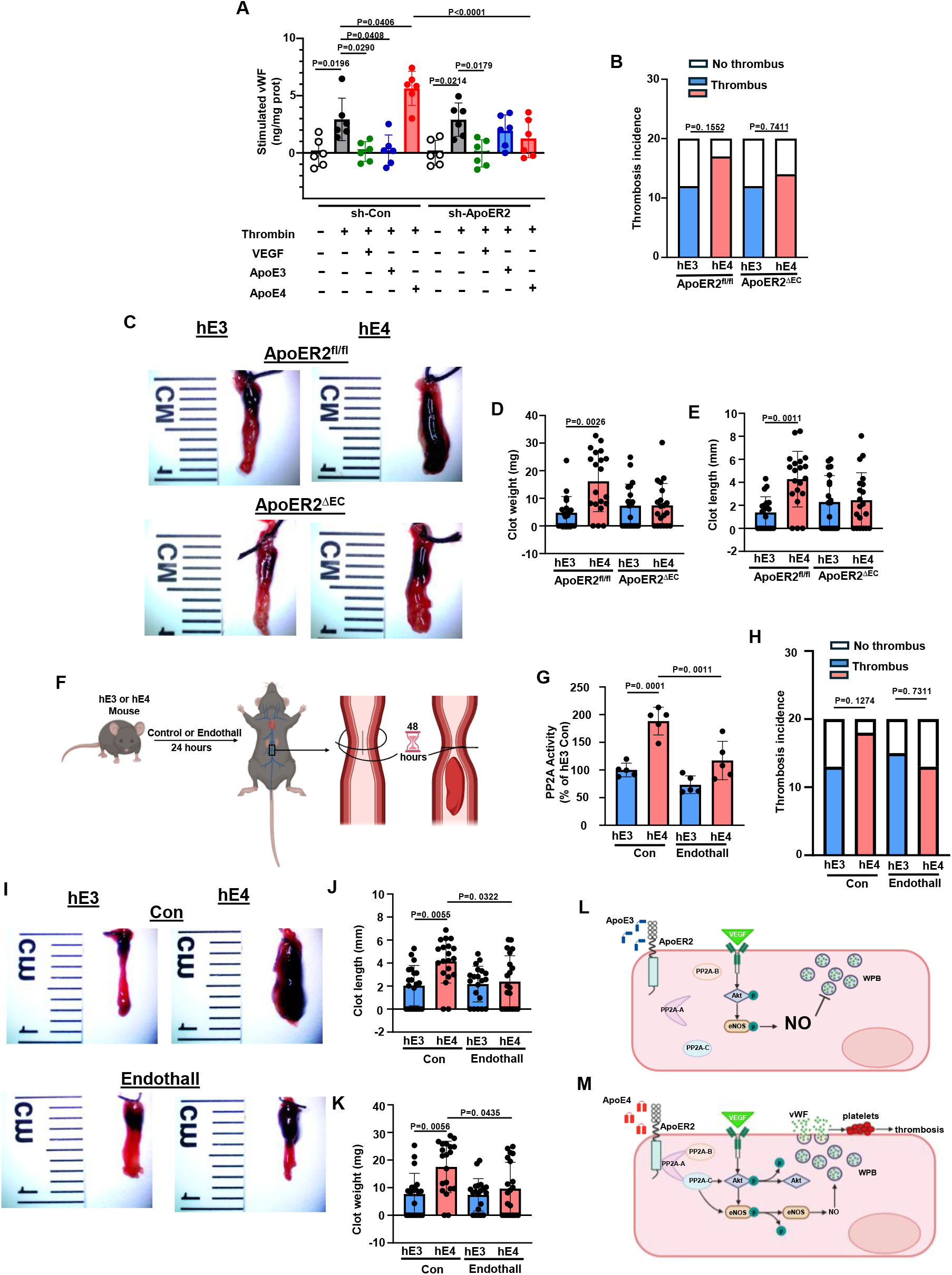
Endothelial cell ApoER2 and PP2A mediate ApoE4 promotion of thrombosis. **A**, To first test effects of ApoE3 and ApoE4 on mouse endothelial cells, vWF secretion by MAEC was measured in the absence or presence of thrombin (3U/ml) plus/minus ApoE3 or ApoE4 (15ug/ml) after control shRNA or shRNA knockdown of ApoER2. **B-E**, The partial flow restriction model of thrombosis in the inferior vena cava (IVC) was performed in male hE3 and hE4 mice with normal endothelial cell ApoER2 expression (ApoER2^fl/fl^) or deficient in endothelial ApoER2 (ApoER2^ΔEC^). The incidence of thrombus formation was evaluated (**B**). Representative images of the thrombus in the dissected IVC are shown in **C**, and clot weight (**D**) and length (**E**) were quantified. N=20 per group. **F-K,** The IVC model of thrombosis was performed in hE3 and hE4 mice treated with the PP2A inhibitor endothall (10 µmol/kg body weight daily) or the control compound 1,4-dimethyl-endothall (**F**). **G**, In a related experiment the IVC were harvested 24h after treatment in the absence of ligation, and PP2A activity was quantified. N=6/group. **H**, The incidence of thrombus formation was evaluated, with N=20/group. **I,** Representative images of the thrombus in the dissected IVC are shown. **J, K**, clot weight (**J**) and length (**K**) were quantified. Values are mean±SEM. Data normality was evaluated by Shapiro-Wild tests. Incidence of thrombosis in the IVC model was compared by Chi-square test (**B, H**). Data were normally distributed and analysis of variance (ANOVA) with Tukey’s post-hoc testing was performed in **A**. In the instances in which normality testing failed, the four groups were compared using by non-parametric ANOVA with Kruskal—Wallis and Dunn’s post-hoc testing (**D, E, G, J, K**). **L, M**, In contrast to events that occur in response to ApoE3 (**L**), the ApoE4 and ApoER2 tandem recruits and activates PP2A, causing dephosphorylation and antagonism of eNOS and NO production, leading to enhanced secretion of vWF and exaggerated thrombosis (**M**).

To implicate PP2A and concurrently test a pharmacological intervention in ApoE4-induced thrombosis in vivo, the effects of the PP2A inhibitor Endothall were tested (Figure 6F). The quantification of PP2A activity in the IVC of hE3 and hE4 mice first showed that activity is 2-fold higher in hE4 (Figure 6G), paralleling the findings in aorta (Figure 4E). Endothall treatment lowered PP2A activity in the hE4 IVC to levels comparable to those found in hE3 IVC. In the IVC model of thrombosis, Endothall blunted the exaggerated thrombus formation in hE4 (Figure 6H-K), fully attenuating both thrombus weight and length. These findings reveal both that elevated PP2A activity underlies the enhanced thrombus formation instigated by ApoE4 in vivo, and that it can be targeted therapeutically.

## DISCUSSION

With possible impact on 25% of the population, *APOE4* is a common genetic risk factor for CVD, and the basis of the increased risk entails mechanisms in addition to alterations in circulating lipids. Whereas it has been demonstrated that ApoE4 increases atherosclerosis severity, whether other major drivers of CVD events are influenced by ApoE4 is unknown. Here we perform GWAS analysis that reveals that *APOE4* raises the risk of VTE, a readily apparent clinical thrombotic disorder. Causality is demonstrated by finding that humanized ApoE4 mice (hE4) have both greater microvascular and venous thrombosis than humanized ApoE3 mice (hE3). The mechanistic underpinnings are then identified, with implication of endothelial cell ApoER2 followed by interactome interrogation revealing a key role for the phosphatase PP2A (Figure 6L,M). In contrast to events that occur in response to ApoE3, the ApoE4 and ApoER2 tandem recruits and activates PP2A, causing dephosphorylation and antagonism of eNOS and NO production, leading to enhanced secretion of vWF and exaggerated thrombosis. In vivo experiments further show that the ApoE4 promotion of thrombosis is exclusively an endothelial cell process, and one that may possibly be amenable to intervention. The newly-identified mechanisms may serve as therapeutic targets for the treatment or prevention of both specific thrombotic disorders like VTE and acute CVD events such as myocardial infarction and stroke in a substantial proportion of individuals.

It has been previously observed that *APOE4* impacts the risk of CVD. In a meta-analysis of 48 CHD studies with a total of 15,492 cases and 32,965 controls, the *APOE4* allele was associated with a 42% greater risk of CVD^50^. In the Baltimore Longitudinal Study of Aging it was found that the *APOE4* genotype is a strong independent risk factor for coronary events in men independent of total cholesterol levels^8^. In a longitudinal study of 258 individuals with similar lipid status on statins, atherosclerosis-related vascular complications occurred more frequently in subjects with *APOE4*^14^. We directly assessed the relationship between *APOE4* and thrombosis by probing a VTE GWAS comprised of multiple independent cohorts with 6,355 VTE cases and 161,416 controls^51^. In the combined GWAS, which surpasses by over 20-fold the number of cases in two prior single cohort studies of *APOE4* and VTE^52,53^, the disorder was more frequent in subjects harboring the variant, clearly demonstrating the association between *APOE4* and thrombosis.

To directly test if and how the *APOE4* genotype impacts thrombosis, we employed two complementary models. Microvascular thrombosis was studied using intravital microscopy in the mesenteric microcirculation and the application of ferric chloride to induce free radical endothelial cell activation. Ferric chloride is the most commonly used method to induce vascular injury to initiate thrombosis^54^. Maximum thrombus size was greater and time to occlusion was shortened in both arterioles and venules in hE4 mice compared to hE3 mice. To model VTE in humans, the ligature stenosis model in the IVC was employed. This model is favored because it lacks an endothelial cell damage intervention and the thrombus forms under existing circulation, and it displays the variability that is seen in humans with VTE^55,56^. Clot weight and length were increased in male and female hE4 mice compared to hE3 mice, paralleling the findings in the microvasculature, and doing so with an entirely different thrombosis initiation method. Observing equal plasma total cholesterol, triglyceride, LDL and HDL levels in hE3 and hE4 mice, the difference in thrombosis severity is not explained by possible effects of ApoE4 on circulating lipids.

Seeking to determine how ApoE4 promotes thrombosis, knowing the critical role of vWF in thrombosis initiation and that endothelial cells are the primary source of vWF influencing hemostasis^46,57^, we evaluated the effect of ApoE3 and ApoE4 on vWF secretion by human endothelial cells. Whereas vWF secretion was attenuated by ApoE3, it was enhanced by ApoE4, and loss-of-function experiments targeting various LDL receptor family members expressed in endothelial cells showed that both responses are mediated by ApoER2. We have previously implicated endothelial cell ApoER2 in the prothrombotic diathesis that characterizes the antiphospholipid syndrome, and in pro-atherogenic actions of Reelin on endothelium^36,58,59^. Recognizing that a major mode of regulation of vWF secretion via Weibel-Palade body (WPB) exocytosis is the modulation of N-ethylmaleimide-sensitive factor (NSF) activity by NO, we determined how ApoE3 and ApoE4 impact NO production and eNOS activation. In contrast to ApoE3, ApoE4 attenuated NO generation in an ApoER2-dependent manner due to a blunting of eNOS activating phosphorylation, and ApoE4 stimulation of vWF secretion was prevented by an NO donor. Thus, in contrast to ApoA3, ApoE4 promotes endothelial cell vWF secretion by negating its attenuation by NO.

The underpinnings of the differential effects of ApoE3 and ApoE4 were then interrogated by studying the interactome of ApoER2 in endothelial cells, finding that it is dynamically modulated by ApoE3 and ApoE4. Seeking an explanation for ApoE4 antagonism of eNOS and the resulting enhancement of vWF secretion, we noted that ApoE3 and ApoE4 cause differential recruitment of members of the PP2A complex and we confirmed the mass spectrometry findings by co-immunoprecipitation. PP2A is a trimeric protein complex in which a core dimer formed between the scaffolding A subunit (PPP2R1A or PPP2R1B) and the catalytic C subunit (PPP2CA or PPP2CB) is associated with one of the many B subunits that facilitate and direct the interaction of the trimer with substrate proteins^48^. Whereas in co-immunoprecipitation ApoE3 and ApoE4 both caused PP2A-A recruitment to ApoER2, PP2A-C was uniquely recruited only in response to ApoE4, which enhanced PP2A phosphatase activity to cause both Akt and eNOS dephosphorylation. Measurements of PP2A activity in mouse aorta further revealed that the ApoE4-induced stimulation occurs readily in vivo. Interestingly, we previously demonstrated participation of endothelial cell PP2A in ApoER2-dependent thrombotic actions of antiphospholipid antibodies^36^. In the current work we have identified a new, robust, genetically-driven activator of PP2A in endothelial cells. Since PP2A modulates endothelial cell barrier function^60–62^, angiogenesis^63^ and apoptosis^64,65^, and it promotes endothelial-to-mesenchymal transition^66^, there may be multiple pathogenic consequences of ApoE4 activation of PP2A in endothelium beyond thrombosis to now consider.

The human relevance of the prothrombotic mechanisms of action of endothelial ApoER2 and PP2A revealed in cell culture and in mice was strengthened by demonstrating that the receptor and enzyme subunits are expressed in human arterial and venous endothelium. Interestingly, in the great saphenous vein ApoER2, encoded by LRP8, is primarily expressed in valvular endothelium as well as being found in venous endothelium. The valve pocket sinus is the most common site of initiation of venous thrombosis, and individuals with more valves have a greater frequency of DVT^67^. And a query of transcriptomes in the mouse IVC venous thrombosis model showed that the endothelial cell mechanisms driving ApoE4 promotion of thrombosis may actually be amplified in the setting of thrombosis. How ApoER2 and PP2A expression are regulated in endothelium represents an additional knowledge gap warranting future interrogation.

ApoE3 and ApoE4, are distinguished by a single point variation at position 112, with ApoE3 having Cys112 and ApoE4 having Arg 112. The presence of Arg112 in ApoE4 exposes a side chain of Arg61, leading to salt-bridge interaction between Arg61 and Glu255 and C-terminal to N-terminal domain folding that underlies its known altered functions compared with ApoE3 ^3–5^. Since the two isoforms have similar high-affinity binding to ApoER2^68^, it remains to be determined how ApoER2-driven mechanisms in endothelium are differentially mediated by ApoE3 and ApoE4. In neurons, compared to ApoE3, ApoE4 attenuates the cell surface recycling of ApoER2 by causing aberrant trafficking in the early endosome^68^. If a similar mechanism occurs in endothelial cells, the altered subcellular localization of ApoER2 may lead to the varied interactomes that we have demonstrated with ApoE3 versus ApoE4, with the latter causing the unique complexing of the three subunits of PP2A and its activation. Prompted by the present findings, the possible role of abnormal ApoER2 trafficking in ApoE4 actions on endothelium, including the participation of the acidic environment in the endosome as implicated in neurons^68^, can be pursued in futures studies.

Along with increasing the risk of CAD^7–10^ and stroke^15,16^, *APOE4* is recognized as the strongest genetic risk factor for late-onset Alzheimer’s disease (AD)^69^. Relevant to the present work, there are numerous forms of evidence that thrombosis contributes to AD pathogenesis. Post-mortem studies of AD patients have revealed increased platelet deposition, thrombin and prothrombin in the CNS vasculature^70,71^, circulating coagulation markers including vWF are increased ante-mortem in AD^72^, and in a number of trials antithrombotic agents have yielded functional improvements in AD patients and in subjects with other neurodegenerative disorders^73^. There is evidence of neurovascular thrombosis and intravascular fibrin(ogen) deposition in mouse models of AD^72,74^, and in the Tg6799 mouse model of AD, treatment with the fibrinogen inhibitor RU-505 blunted the prothrombotic phenotype and improved cognitive function^75^. When interrogating the expression of ApoER2 (LRP8) in endothelial cells, we discovered that it is particularly abundant in human CNS endothelial cells. Thus, our discovered prothrombotic features of the ApoE4 and endothelial cell ApoER2 tandem may be operative in the CNS, and they may be highly relevant to the detrimental impact of ApoE4 on AD occurrence and severity. The finding that LRP8 is most abundant in the blood-brain barrier endothelium has additional significance because ApoE4-induced blood-brain barrier disruption may contribute to ApoE4-associated cognitive decline independent of AD pathology^76^.

The COVID-19 pandemic may have been a natural experiment in which the influence of ApoE4 on hemostasis was apparent. Microangiopathy with microvascular thrombosis was a major pathogenic as well as clinical feature of severe COVID-19 infection, with excessive endothelial cell activation causing marked changes in hemostasis that led to venous and arterial thrombosis in the lung and in other organs^77^. Consistent with this, it was observed that whereas circulating vWF was not raised in individuals with mild disease, it was increased 2.7-fold compared to normal levels in severe COVID-19 patients^78^. In a study early in the pandemic, compared to individuals homozygous for *APOE3*, in subjects homozygous for *APOE4* the risk of severe COVID-19 illness was more than doubled^79^. This was borne out in a later meta-analysis of nine independent studies in which *APOE4* carriers compared to *APOE3* carriers had an increased risk of severe disease (OR 1.85, CI 1.50-2.28)^80^. The promotion of thrombosis by ApoE4 and ApoER2 may have contributed to the greater likelihood of disease progression upon COVID-19 infection in individuals harboring the variant.

In the realm of immunotherapy for cancer it has been observed that PP2A may be a therapeutic target. PP2A inhibition has been found to enhance anticancer immunity, with one example being the enhancement of the efficacy of CAR-T cell therapy against glioblastoma in a murine model^81^. In addition, patients with tumors with inactivating somatic mutations in PPP2R1A, the gene encoding the most common A subunit in the PP2A complex, had improved survival in a number of immune checkpoint blockade-treated patient cohorts across multiple cancer types^82^. The further development of PP2A inhibition therapies in the context of cancer may lead to innovations that will benefit the vascular health of individuals harboring ApoE4. Springboarding from the discoveries reported here, future additional insights into the ApoE4-ApoER2 partnership in endothelial cells may also increase the feasibility of breaking the link between ApoE4 and both primary thrombotic disorders like VTE and acute CVD events such as myocardial infarction and stroke in 25% of the population.

## Acknowledgements

The authors thank the UT Southwestern Proteomics Core for assistance with proteomics analyses.

## Author Contributions

Dr. Sun conducted the mouse inferior vena cava ligation studies of venous thrombosis, the quantification of serum lipids, the studies of endothelial cell vWF secretion, immunoblotting to evaluate shRNA-based gene excision in cell culture and protein phosphorylation, co-immunoprecipitation, measurements of NO production through quantification of nitrate and nitrite release, the quantification of PP2A enzyme activity in cell culture and in tissues, and contributed to manuscript writing and figure production. Dr. Sacharidou performed the intravital microscopy experiments interrogating microvascular thrombosis and related data analyses, conducted PP2A enzyme activity assays, and participated in manuscript writing and figure generation. Dr. Chen performed the GWAS analyses and the interrogation of publicly-available single cell RNAseq data sets for gene expression in vascular cells. Dr. Chen also contributed to manuscript writing and figure generation. Dr. Lemoff conducted the liquid chromatography and tandem mass spectrometry, and contributed to manuscript preparation. Drs. Gaddam and Lella Rao provided instruction in the mouse inferior vena cava ligation model of venous thrombosis, and contributed to manuscript writing. Dr. Xu provided supervision of the GWAS analyses and the interrogation of single cell RNAseq data sets, and participated in manuscript writing and figure generation. Dr. Mineo contributed to study design, data analyses and interpretation, and manuscript writing and figure generation. Dr. Shaul conceived and designed studies, interpreted data, and drafted and revised the text and figures.

## Data Availability

The mass spectrometry proteomics data have been deposited to the ProteomeXchange Consortium (http://proteomecentral.proteomexchange.org) via the MassIVE partner repository with the dataset identified PXD080691.

## Supplemental Material

Detailed Methods

Supplemental Tables

Supplemental Figures and Figure Legends

Legends for Video files

Video Files

## Abbreviations

AD: Alzheimer’s disease
ANOVA: one-way analysis of variance
ApoE: apolipoprotein E
ApoER2: apolipoprotein E receptor 2
CAD: coronary artery disease
CVD: cardiovascular disease
DVT: deep vein thrombosis
eNOS: endothelial cell NO synthase
HAEC: human aortic endothelial cells
hE3: humanized *APOE3*
hE4: humanized *APOE4*
IVC: inferior vena cava
LC/MS-MS: Liquid Chromatography/ Tandem Mass Spectrometry
MAEC: mouse aortic endothelial cells
NO: nitric oxide
NSF: N-ethylmaleimide-sensitive factor
PE: pulmonary embolism
PP2A: serine/threonine protein phosphatase 2A
SNARE: soluble NSF attachment protein receptor
VTE: venous thromboembolism
vWF: von Willibrand Factor
WPB: Weibel-Palade bodies

