## Supplemental Materials for "ApoE4 Promotes Thrombosis via Endothelial Cell ApoER2 and PP2A Activation"

#### Detailed Methods

GWAS Analyses

To evaluate the association between *APOE4* and venous thromboembolism (VTE), raw SNP array genotype data were downloaded from a venous thromboembolism (VTE) GWAS dataset reported by Munsch et al.[1] and used for a focused regional association analysis centered on the *APOE* locus to assess statistical associations between SNP genotypes and VTE susceptibility. After applying standard genotype- and sample-level quality control procedures, VTE case-control status was tested against each SNP genotype under an additive genetic model using logistic regression, and rs429358 was interpreted in the context of surrounding linkage disequilibrium. The regression coefficient is interpreted as the log odds ratio (OR), with OR = exp(beta); when only an OR is available, beta is calculated as ln(OR). When a standard error can be computed, 95% confidence intervals (CI) are calculated as exp[beta +/- 1.96 x standard error (SE)] for ORs and beta +/- 1.96 x SE. When a 95% confidence interval can be computed but the SE is not, the SE is derived as [ln(upper CI) - ln(lower CI)] / (2 x 1.96) for ORs or (upper CI - lower CI) / (2 x 1.96). The two-sided Wald test is used to calculate P value, with z = beta/SE and P = 2 x [1 - Phi(|z|)]. All associations are harmonized to the APOE4-defining allele at rs429358 (C allele on the plus strand): associations for the opposite allele are inverted by multiplying beta by -1 or by taking the reciprocal of the odds ratio, while the two-sided P value is left unchanged. The effective sample size is calculated as

4 / (1/Ncases + 1/Ncontrols) when case and control counts are available. For comparison with the interrogation of APOE4 and VTE, to evaluate the association between APOE4 and other cardiovascular phenotypes, additional GWAS cohorts were required. Separate analyses were necessary because typical GWAS datasets evaluating cardiovascular parameters lack data on VTE. For the additional cardiovascular phenotypes, rs429358/APOE4 associations were examined based on the same procedure as detailed above. For these additional cardiovascular phenotypes, because the contributing datasets differed in ancestry composition, phenotype ascertainment, genotyping or imputation platform, covariate adjustment, and association-analysis pipeline, estimates were summarized descriptively in the sun plot (Figure 1) and were not pooled in a formal cross-phenotype meta-analysis.

Animal Models

We previously demonstrated effective and selective excision of ApoER2 from endothelium in ApoER2$^{\Delta EC}$ mice [2,3]. In select experiments hE3 and hE4 mice were treated daily by intraperitoneal injection with the PP2A inhibitor Endothall (10 μmol/kg body weight, Calbiochem, Cat. No. CAS 145-73-3) or the control compound 1,4-dimethyl-endothall (Santa Cruz, Cat. No. CAS 109282-27-1)[2] for 1 to 3 days. The mice were housed at 23°C with light cycles of 12 h of light beginning at 6:00am and 12 h of dark beginning at 6:00pm, humidity was 30-70%, and food and $H_2O$ were provided ad libitum. Circulating lipids were evaluated by quantification of plasma total cholesterol, total triglycerides, LDL and HDL[4]. All animal experiments were approved by the Institutional

Animal Care and Utilization Committees at the University of Texas Southwestern Medical Center.

Intravital Microscopy

The intravital microscopy have been previously reported[2]. Briefly, mice (males, 4-6 weeks old) were IV injected with fluorescence-labeled anti-mouse GPIbβ antibody (0.1ug/g body weight, Emfret Analytics cat. no. X488), the mesenteric microcirculation was exteriorized and thrombus formation was induced by 10% ferric chloride solution placed on Whatman filter paper for 45 sec. Thrombus formation in 80–100 μm arterioles and accompanying venules was documented by capturing images (X200 magnification) every 1 second for 25 min with a QuantEm 512S camera attached to a NIKON Eclipse Ti microscope with NIS Element image capturing software. The largest thrombus formed within 7 min in arterioles and within 3 min in venules, and time to full occlusion were determined in five vessels per mouse, and the mean value per mouse was calculated.

IVC Model of Thrombosis

A partial flow restriction (stenosis) model of venous thrombosis in the inferior vena cava (IVC) was used to provide a complementary phenotyping of thrombosis[5-7]. At 8 to 10 weeks of age male or female mice were anesthetized with isoflurane at 2% and an oxygen flow rate of 200 ml/min and placed in a supine position on a PhysioSuite pad to maintain body temperature, which was monitored with a temperature probe. After laparotomy, the intestines were exteriorized and sterile saline was applied during the entire procedure to prevent drying. After gentle separation from the aorta, the IVC was ligated with a 7-0 polypropylene suture immediately inferior to the renal veins, placing a 30-gauge needle on the exterior of the IVC to invoke incomplete ligation, and the needle was then immediately removed. This procedure decreases the vascular lumen area to 10% of the intact vessel and allows for regimented flow restriction without endothelial injury. All IVC side branches, typically two to four in number, were occluded by cauterization. After surgery, the peritoneum and skin were closed using 5-0 nylon suture and the mouse was allowed to recover. Mice were euthanized after 48 hours, and the IVC inferior to the suture and the thrombi that had developed were harvested. The incidence of thrombosis and thrombus length and weight were determined. In additional experiments the IVC was harvested without manipulation for the measurement of PP2A enzymatic activity.

<u>Cell Culture Models</u>

Human aortic endothelial cells (HAEC) were purchased from Lonza and maintained in EGM-2 and Endothelial Growth Medium with added growth factors (Lonza). Mouse aortic endothelial cells (MAEC) were obtained from iXCells Biotechnologies and maintained in Complete Mouse Endothelial Cell Medium (Cell Biologics, Cat. No. M1168). Cells were used within 3-5 passages. shRNA was employed to knockdown LDLR, VLDLR, LRP1, ApoER2 and PP2A-A (PPP2R1) in HAEC, and ApoER2 in MAEC, using lentiviral constructs generated by VectorBuilder. All experiments included an shRNA control. Effective gene silencing was evaluated by immunoblotting using anti-LDLR (1:1000, R&D systems, cat. no. AF2148), anti-VLDLR (1:1000, R&D Systems, cat. no. AF2148), anti-LRP1 (1:1000, Abcam, cat. no. ab92544), anti-ApoER2 (1:1000, Abcam, cat. no. ab108208), anti-PP2A-A (1:1000, Cell Signaling Technology, cat. no. 2039) and anti-actin (1:2000, Santa Cruz Biotechnology, catalog no. sc47778). Experiments were performed 48h following gene knock-down.

vWF Secretion

Cells plated in 6-well plates were incubated with vehicle or thrombin (3U/ml, Sigma-Aldrich, cat. no. T1063), and vWF accumulated in the media over 2h was measured by ELISA (Invitrogen, catalog no. EHVWF). Additional treatments included VEGF (100ng/ml, Sigma, catalog no. SRP3182), ApoE3 or ApoE4 (15ug/ml, Peprotech, cat. nos. 350-02 and 350-04). In select studies the cells were additionally treated with the NO donor spermine NONOate (1uM, Cayman Chemical, cat. no. 82150) or the control parent compound spermine (Cayman Chemical, cat. no. 18041)[8,9]. To further evaluate the role of PP2A, additional studies were performed in cells treated with the PP2A inhibitor Endothall or the control compound 1,4-dimethyl endothall (1uM)[2].

eNOS Phosphorylation and Activation

eNOS activity was assessed in HAEC using two approaches. NO production was determined by the measurement of total nitrate and nitrite in the media following 15min incubation in the absence or presence of VEGF (100ng/ml) plus or minus the addition of the NOS antagonist nitro-L-arginine-methyl ester (L-NAME, 2mM). Activating phosphorylation of eNOS was evaluated by immunoblotting for phosphorylated eNOS-Ser1177 (1:1000, Cell Signaling Technology, catalog no. 9570) and total eNOS (1:1000, Cell Signaling Technology, catalog no. 9572). Akt activation was also evaluated by immunoblotting for phosphorylated Akt-Ser473 (1:1000, Cell Signaling Technology, catalog no. 9271S) and total Akt (1:1000, Cell Signaling Technology, catalog no. 9272S).

Immunoprecipitation and Immunoblot Analysis

Previously-described methods[10] were modestly modified to assess protein co-immunoprecipitation after cell incubation with vehicle versus ApoE3 or ApoE4 (15ug/ml) for 15 min. Immunoprecipitation was done with mock IgG control or ApoER2 antibody (2.5ug, Abcam #ab108208). Immunoblot analyses, which included samples of cell lysates representing the co-immunoprecipitation inputs, were performed using antibodies against ApoER2 and subunits of the protein phosphatase PP2A. PP2A is a trimeric protein complex in which a core dimer formed between the scaffolding A subunit (PPP2R1A or PPP2R1B) and the catalytic C subunit (PPP2CA or PPP2CB) is associated with one of the many B subunits that facilitate and direct the interaction of the trimer with substrate proteins[11]. PP2A-A (1:1000, Cell Signaling Technology, catalog no. 2039) and PP2A-C (1:1000, Cell Signaling Technology, catalog no. 2038) were detected. Additional immunoblotting was performed detecting LDLR, VLDLR, LRP1 and actin.

Liquid Chromatography/ Tandem Mass Spectrometry (LC/MS-MS)

To evaluate the endothelial cell ApoER2 interactome and its dynamic changes in response to ApoE3 versus ApoE4, HAEC were treated with vehicle, ApoE3 or ApoE4 (15ug/ml for 15 min), ApoER2 was immunoprecipitated, and the associated proteins were evaluated by liquid chromatography/tandem mass spectrometry (LC/MS-MS)[49]. Three biological replicates were used for each condition. Following protein separation by sodium dodecyl sulfate poly-acrylamide gel electrophoresis, gel samples were digested overnight with trypsin (Pierce) followed by reduction and alkylation with dithiothreitol and iodoacetamide (Sigma-Aldrich). After solid-phase extraction cleanup with Oasis HLB plates (Waters), the samples were injected onto a 75 um i.d., 15-cm long EasySpray column (Thermo) and eluted with a gradient from 0-28% buffer B over 90 min. Buffer A contained 2% (v/v) ACN and 0.1% formic acid in water, and buffer B contained 80% (v/v) ACN, 10% (v/v) trifluoroethanol, and 0.1% formic acid in water. Samples were injected onto a Thermo Q Exactive HF mass spectrometer coupled to a Thermo Vanquish Neo nano liquid chromatography system. The mass spectrometer operated in positive ion mode with a source voltage of 2.5 kV and an ion transfer tube temperature of 300 °C. MS scans were acquired at 120,000 resolution in the Orbitrap and up to 20 MS/MS spectra were obtained in the ion trap for each full spectrum acquired using higher-energy collisional dissociation (HCD) for ions with charges 2-8. Dynamic exclusion was set for 20 s after an ion was selected for fragmentation. Raw MS data files were analyzed using Proteome Discoverer v.3.3 (Thermo), with peptide identification performed using Sequest HT searching against the human reviewed protein database from UniProt. Fragment and precursor tolerances of 10 ppm and 0.02 Da were specified, and three missed cleavages were allowed. Carbamidomethylation of Cys was set as a fixed modification, with oxidation of Met set as a variable modification. Peaks were detected using the Minora Feature Detector within Proteome Discoverer. The proteins were chosen based upon the FDR Confidence (<1% false discovery rate) and number of Peptide Spectrum Matches (#PSMs), or the number of spectra assigned to peptides that contributed to the inference of the protein (#PSMs>2). Protein abundance values were calculated as the sum of the peak intensities for each peptide identified for that protein. The abundance values were adjusted such that the sums of the protein abundances for each sample are equal, to correct for differences in protein loading amount. Protein abundance for three biological replicates was averaged, the abundance with ApoE3 treatment versus vehicle and the abundance with ApoE4 treatment versus vehicle were calculated, and the results were tabulated as proteins more abundant with ApoE3 versus ApoE4 or more abundant with ApoE4 versus ApoE3.

PP2A Activity

PP2A phosphatase activity was evaluated using PP2A immunoprecipitation and a phosphatase assay kit[2]. Cultured endothelial cells were incubated for 90 min in the presence of vehicle, ApoE3 or ApoE4 (15 ug/ml), and C2-ceramide (30 μM) served as positive control. Lysates were generated and immunoprecipitation was performed with anti-PP2A-C antibody immobilized on agarose beads for 2h. Immunoprecipitated PP2A was incubated with phosphorylated peptide substrate for 10 min at 30°C. The reaction was terminated by the addition of malachite green phosphate detection reagent, and the amount of free phosphate released was quantified by measuring absorbance at 650 nm using a microplate reader. Phosphate concentrations were calculated from a phosphate standard curve generated in parallel, and PP2A phosphatase activity was expressed relative to vehicle control. Using a parallel approach PP2A activity was quantified in mouse aorta or IVC.

Vascular Cell Transcriptome Analyses

Publicly available single cell RNA-seq datasets relevant for vascular biology and thrombosis were interrogated to determine the distribution of expression of LRP8 and PP2A subunit genes in vascular cells. We collected single cell RNA-seq data from Barnett et al (2024)[12], Sun et al (2022)[13] and DeRoo et al (2023)[14]. The datasets are provided in different formats, and as a result a variety of processing approaches were required. In brief, for a dataset with only raw fastq files available, the 10X cellranger pipeline was run to obtain the raw count matrix for each sample. For a dataset provided as a .h5ad file, the file was converted to .rds format using the sceasy R package. In the case of a dataset provided as a .h5 file, the Read10X_h5() function from Seurat R Package was used to import the raw count matrix. For those datasets provided as raw count matrix, the count matrix was imported using the Seurat R package. For datasets which provide cell-type annotation, the original annotation was used. When annotation did not accompany raw count files, preprocessing and integration were performed using Seurat package. Following the integration of all the samples into one dataset, unsupervised clustering was performed using the FindCluster() function of the Seurat R package, the same number of clusters as shown in the original work were generated, and dotplots of marker gene expression aided manual annotation.

**Supplemental Figures**

### Figure S1

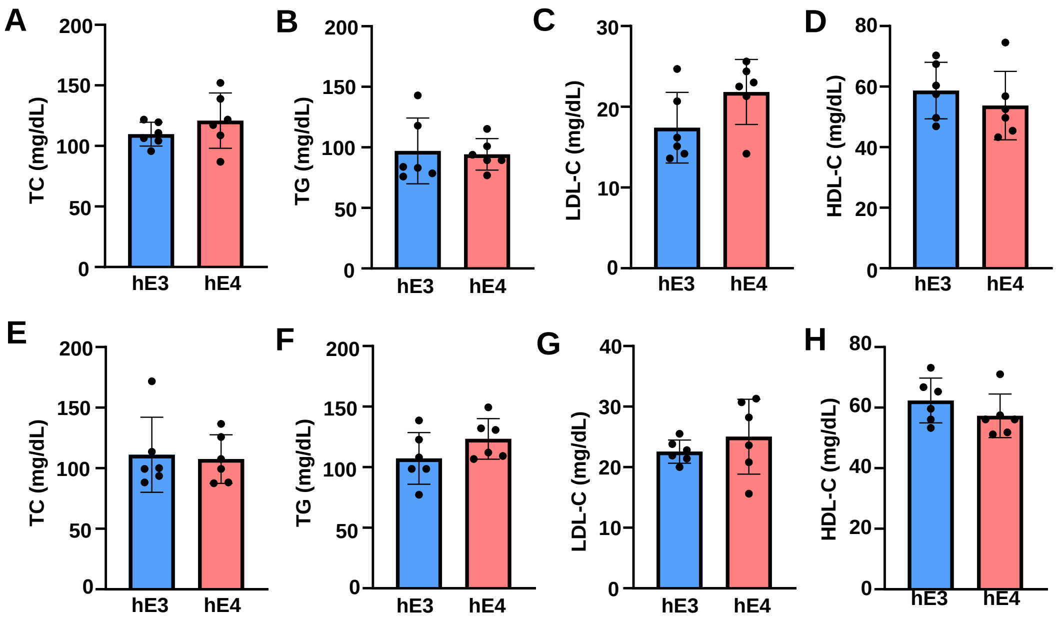


**Figure S1. ApoE4 in mice does not impact circulating lipids.** Following weaning, male (**A-D**) and female (**E-H**) hE3 and hE4 mice were placed on standard chow and at age 8-10 weeks levels of plasma total cholesterol (TC; **A,E**), triglyceride (TG; **B,F**), LDL cholesterol (LDL-C; **C,G**) and HDL cholesterol (HDL-C; **D,H**) were measured (n=6 per group). Values are mean±SEM. Data normality was confirmed by Shapiro-Wilk tests, and comparisons between two groups were evaluated by two-sided, unpaired Welch's t test.

### Figure S2

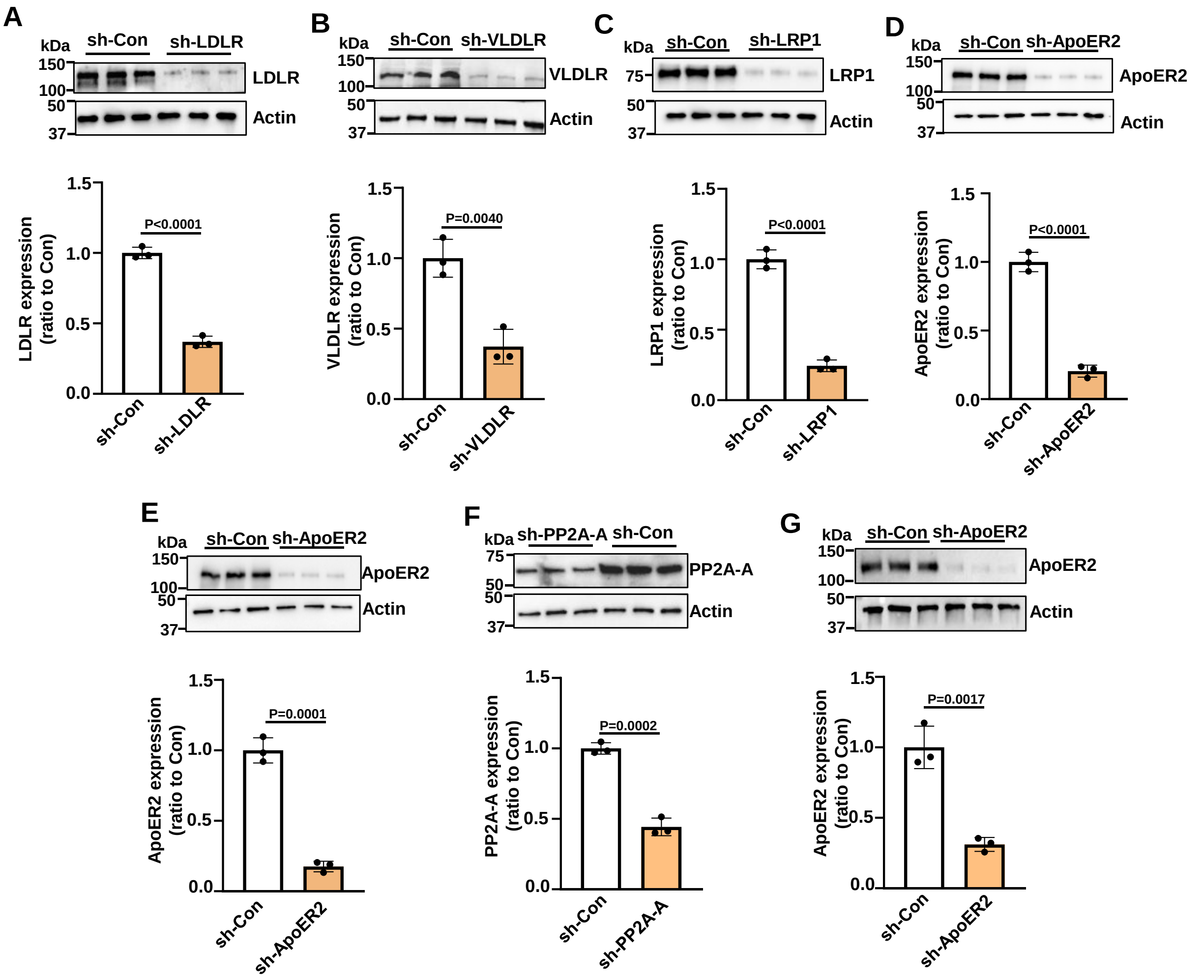


**Figure S2. ApoE receptor and PP2A-A knockdown in cultured endothelial cells. A-E,** ApoE receptor expression in HAEC was evaluated by immunoblotting following control shRNA or shRNA knockdown of LDLR (**A**), VLDLR (**B**), LRP1 (**C**), or ApoER2 (**D, E**). **F**, PP2A-A knockdown in HAEC, and **G**, ApoER2 deletion in MAEC. Upper panel displays immunoblots and quantification is in lower panel. Values are mean±SEM, n=3/group. Groups were compared by two-sided, unpaired Welch's t tests.

### Figure S3

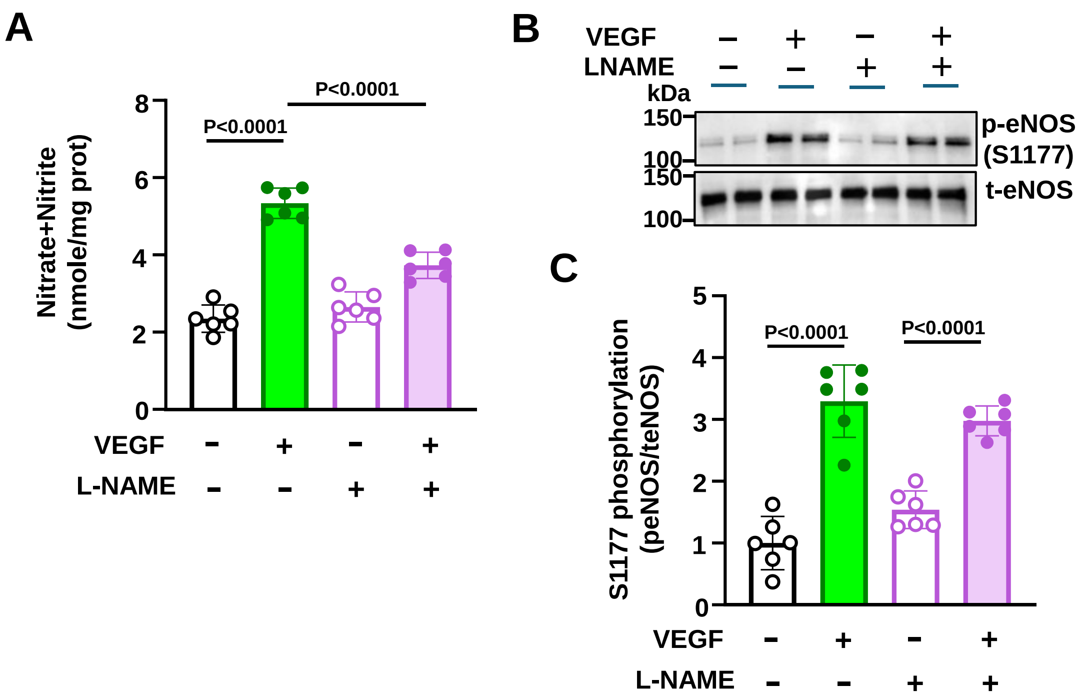


**Figure S3. Establishment of measurement of NO production and eNOS Ser-1177 activating phosphorylation detection in HAEC. A**, Cells were treated with vehicle or VEGF (100ng/ml) in the absence or presence of L-NAME (2mM), and NO production was determined by the measurement of total nitrate and nitrite in the media, n=6/group. **B, C**, Phosphorylated eNOS Ser-1177 and total eNOS were detected by immunoblotting. **B,** displays representative immunoblots with two samples per treatment, and quantification is in **C** (n=6/group). Values are mean±SEM. Datasets were normally distributed, and groups were compared by analysis of variance (ANOVA) with Tukey's post-hoc testing.

**Figure S4**

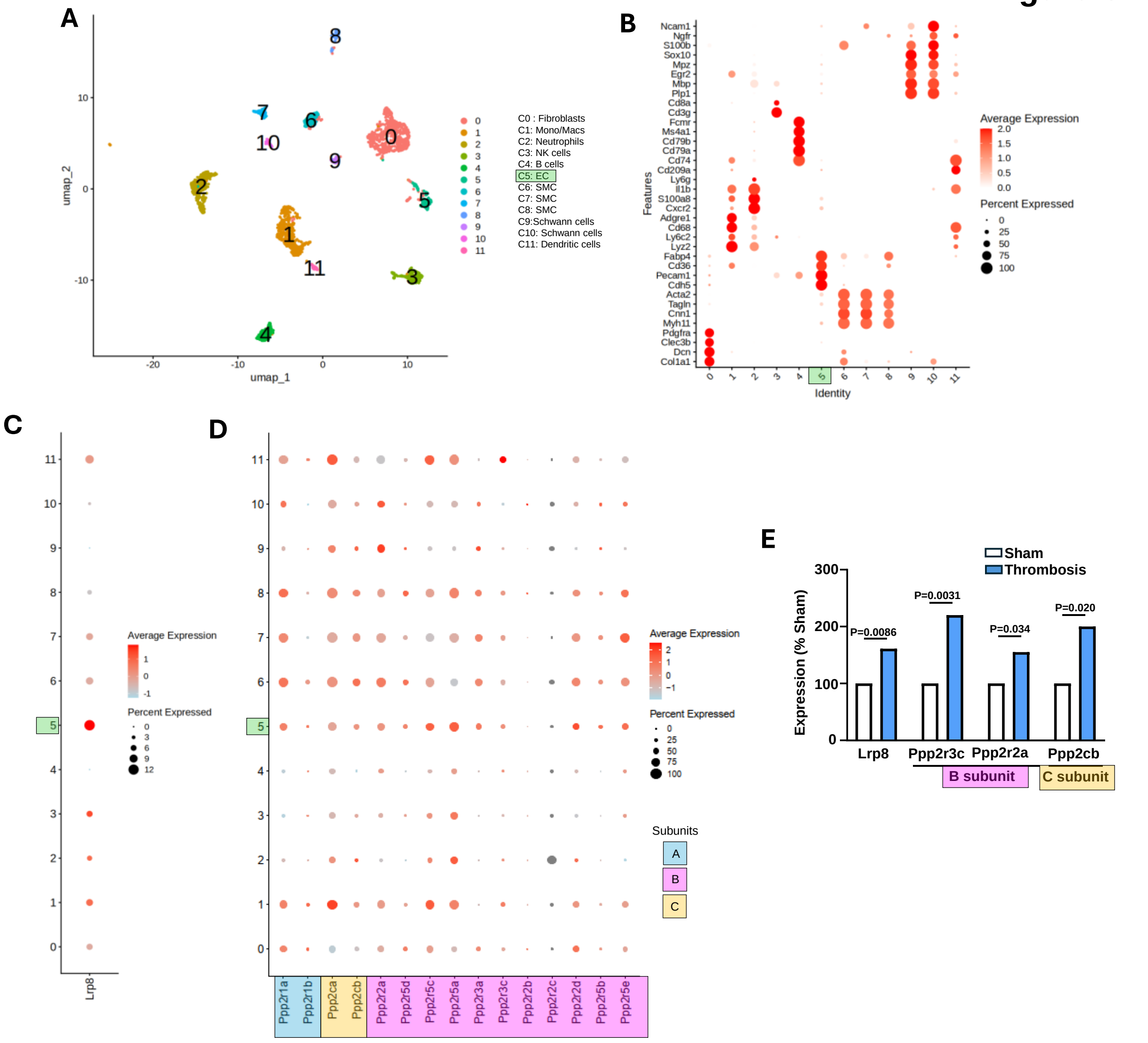


**Figure S4. Endothelial cell ApoER2 (LRP8) and PP2A expression are upregulated in the setting of thrombosis. A, B,** UMAP plot of cell type clusters in mouse inferior vena cava, or IVC (**A**), and marker gene expression summary (**B**). **C**, Dot plot for expression of Lrp8 in various venous cells. **D**, Dot plot for expression of PP2A subunit genes in various venous cells. **E**, Relative expression of Lrp8 and PP2A B subunit and C subunit genes in the IVC endothelium following sham instrumentation versus stenosis-induced thrombosis. Unadjusted P values are shown.

#### Supplemental Tables

**Table S1.** GWAS phenotypes, P values and dataset sizes and sources.

**Table S2.** Proteins associated with ApoER2: those more abundant with ApoE3 compared to ApoE4 treatment, and those more abundant with ApoE4 compared to ApoE3 treatment.

**Table S1**. GWAS phenotypes, P values and dataset sizes and sources

GWAS phenotypes, P values and dataset sizes

| **SNP ID** | **GWAS phenotype in short** | **GWAS phenotype** | **Chromsom** | **locus** | **P-value** | **dataset size** |
|---|---|---|---|---|---|---|
| rs429358 | CAD | description:"Coronary artery disease (CAD)" | 19 | 45411941 | 1.163e-45 | 1360560 |
| rs429358 | AbAA | description:"Abdominal aortic aneurysm" | 19 | 45411941 | 5.406e-39 | 1125328 |
| rs429358 | MI | description:"Myocardial infarction" | 19 | 45411941 | 1.534e-30 | 487368 |
| rs429358 | niHF | description:"Non-ischemic heart failure" | 19 | 45411941 | 1.19E-06 | 162758 |
| rs429358 | CarotidArteryPlaqueBurden | description:"Carotid artery plaque burden" | 19 | 45411941 | 0.000791 | 26807 |
| rs429358 | ChronicPulmoHeartDisease | description:"Chronic pulmonary heart disease" | 19 | 45411941 | 0.00406 | 62680 |
| rs429358 | VenousThromboembolism | description:"Venous Thromboembolism" | 19 | 45411941 | 0.0042 | 168116 |
| rs429358 | MyocardialT1 | description:"Native myocardial T1 time" | 19 | 45411941 | 0.00675 | 86535 |
| rs429358 | LVESVI | description:"Left ventricular end-systolic volume (BSA-indexed)" | 19 | 45411941 | 0.008059 | 113260 |
| rs429358 | LVEDV | description:"Left ventricular end-diastolic volume" | 19 | 45411941 | 0.04793 | 94097 |
| rs429358 | DescAortaStrain | description:"Descending aorta strain" | 19 | 45411941 | 0.052 | 42342 |
| rs429358 | LVESV | description:"Left ventricular end-systolic volume" | 19 | 45411941 | 0.05642 | 94097 |
| rs429358 | LAmax | description:"Indexed left atrial maximum volume" | 19 | 45411941 | 0.058 | 35648 |
| rs429358 | Atherosclerosis | description:"Atherosclerosis" | 19 | 45411941 | 0.0611 | 105887 |
| rs429358 | niHFrEF | description:"Non-ischemic heart failure with reduced ejection fraction" | 19 | 45411941 | 0.06168 | 19901 |
| rs429358 | CongestiveHeartFail | description:"Congestive heart failure" | 19 | 45411941 | 0.075 | 89416 |
| rs429358 | LVESA | description:"Left ventricular end systole anterior wall thickness" | 19 | 45411941 | 0.094 | 41581 |
| rs429358 | HYPERTENSION | description:"Hypertension" | 19 | 45411941 | 0.09439 | 1047140 |
| rs429358 | PulmonaryHT | description:"Pulmonary hypertension" | 19 | 45411941 | 0.09957 | 15941 |
| rs429358 | LAPEF | description:"Left atrial passive emptying fraction" | 19 | 45411941 | 0.1 | 35648 |
| rs429358 | LVmass-EDVratio | description:"Left ventricular mass to end-diastolic volume ratio" | 19 | 45411941 | 0.1 | 16884 |
| rs429358 | HypertrophicCardiomyopathy( | description:"Hypertrophic cardiomyopathy (sarcomere-negative)" | 19 | 45411941 | 0.10408 | 14048 |
| rs429358 | PADnonT2D | description:"Peripheral artery disease in subjects without diabetes" | 19 | 45411941 | 0.11 | 20978 |
| rs429358 | DescAortaDisten | description:"Descending aorta distensibility" | 19 | 45411941 | 0.12 | 42342 |
| rs429358 | LVC | description:"Left ventricular concentricity (left ventricular mass / left ve | 19 | 45411941 | 0.13 | 36083 |
| rs429358 | HypertrophicCardiomyopathy( | description:"Hypertrophic cardiomyopathy (MTAG)" | 19 | 45411941 | 0.13736 | 28106 |
| rs429358 | DescAortaDiam | description:"Descending aorta diameter" | 19 | 45411941 | 0.1457 | 75762 |
| rs429358 | CardiacArrhythmia | description:"Arrhythmia (cardiac)" | 19 | 45411941 | 0.1781 | 160878 |
| rs429358 | VFibFlutter | description:"Ventricular fibrillation and flutter" | 19 | 45411941 | 0.1861 | 4660 |
| rs429358 | DeepVeinThrombosis | description:"Blood clot / pulmonary embolism / deep vein thrombosis ( | 19 | 45411941 | 0.1887 | 49339 |
| rs429358 | AorticStenosis | description:"Calcific aortic stenosis" | 19 | 45411941 | 0.1923 | 53901 |
| rs429358 | ThoracicAA | description:"Thoracic aortic aneurysm" | 19 | 45411941 | 0.1989 | 19646 |
| rs429358 | DPAcm | description:"Short-axis diastolic pulmonary artery (cm)" | 19 | 45411941 | 0.2 | 41135 |

| | | | | | | |
|---|---|---|---|---|---|---|
| rs429358 | Cardiomyopathy | description:"Hypertrophic cardiomyopathy" | 19 | 45411941 | 0.234646 | 21725 |
| rs429358 | SND_INC | description:"Sinus node dysfunction (inclusive)" | 19 | 45411941 | 0.251 | 25410 |
| rs429358 | SystolicHeartFail | description:"Heart failure with reduced EF [Systolic or combined heart f | 19 | 45411941 | 0.253 | 160769 |
| rs429358 | LVEDIL | description:"Left ventricular end diastole inferolateral wall thickness" | 19 | 45411941 | 0.27 | 42194 |
| rs429358 | RVEDV | description:"Right ventricular end-diastolic volume" | 19 | 45411941 | 0.28 | 41135 |
| rs429358 | RVESV | description:"Right ventricular end-systolic volume" | 19 | 45411941 | 0.28 | 41135 |
| rs429358 | AscAortaDisten | description:"Ascending aorta distensibility" | 19 | 45411941 | 0.31 | 42342 |
| rs429358 | PADinT2D | description:"Peripheral artery disease in type 2 diabetes (T2D)" | 19 | 45411941 | 0.32 | 12237 |
| rs429358 | PAcm | description:"Short-axis pulmonary artery (cm)" | 19 | 45411941 | 0.34 | 41135 |
| rs429358 | PAD_eversmk | description:"Peripheral artery disease in ever smokers" | 19 | 45411941 | 0.36 | 22814 |
| rs429358 | RVEFtoLVEF | description:"Right ventricular ejection fraction to left ventricular ejectic | 19 | 45411941 | 0.36 | 41135 |
| rs429358 | NICM | description:"Nonischemic cardiomyopathy" | 19 | 45411941 | 0.3634 | 7230 |
| rs429358 | LAAEF | description:"Left atrial active emptying fraction" | 19 | 45411941 | 0.37 | 35648 |
| rs429358 | SVL | description:"Left ventricular stroke volume" | 19 | 45411941 | 0.3823 | 77177 |
| rs429358 | EssentialHYPERTENSION | description:"Essential hypertension" | 19 | 45411941 | 0.3885 | 459952 |
| rs429358 | LVESAS | description:"Left ventricular end systole anterospetal wall thickness" | 19 | 45411941 | 0.43 | 41770 |
| rs429358 | LVEDA | description:"Left ventricular end diastole anterior wall thickness" | 19 | 45411941 | 0.44 | 41699 |
| rs429358 | CardiacArrest | description:"Cardiac arrest" | 19 | 45411941 | 0.449 | 9476 |
| rs429358 | IVS | description:"Interventricular septum" | 19 | 45411941 | 0.45 | 24598 |
| rs429358 | RVESVtoLVESV | description:"Right ventricular end-systolic volume to left ventricular en | 19 | 45411941 | 0.46 | 41135 |
| rs429358 | RVSV | description:"Right ventricular stroke volume" | 19 | 45411941 | 0.46 | 41135 |
| rs429358 | LVDD | description:"Left ventricular diastolic dimension" | 19 | 45411941 | 0.48 | 24965 |
| rs429358 | LVEDIS | description:"Left ventricular end diastole inferoseptal wall thickness" | 19 | 45411941 | 0.48 | 42098 |
| rs429358 | LVEDAS | description:"Left ventricular end diastole anterospetal wall thickness" | 19 | 45411941 | 0.49 | 41286 |
| rs429358 | LVGLS | description:"Left ventricular global longitudinal strain" | 19 | 45411941 | 0.52 | 35052 |
| rs429358 | PAD | description:"Peripheral artery disease" | 19 | 45411941 | 0.53 | 40017 |
| rs429358 | LVESIS | description:"Left ventricular end systole inferoseptal wall thickness" | 19 | 45411941 | 0.54 | 41984 |
| rs429358 | LVmassIndex | description:"Left ventricular mass index" | 19 | 45411941 | 0.54 | 22691 |
| rs429358 | SupraventricularPremaBeats | description:"Supraventricular premature beats (PheCode 427.61)" | 19 | 45411941 | 0.5426 | 10352 |
| rs429358 | LVEF | description:"Left ventricular ejection fraction" | 19 | 45411941 | 0.5512 | 130183 |
| rs429358 | AscAortaDiam | description:"Ascending aorta diameter" | 19 | 45411941 | 0.5613 | 116897 |
| rs429358 | SCAD | description:"Spontaneous coronary artery dissection" | 19 | 45411941 | 0.5774 | 6357 |
| rs429358 | AVNRT | description:"Atrioventricular nodal reentrant tachycardia (AVNRT)" | 19 | 45411941 | 0.5775 | 9327 |
| rs429358 | RVSVtoLVSV | description:"Right ventricular stroke volume to left ventricular stroke vo | 19 | 45411941 | 0.59 | 41135 |
| rs429358 | Cardiomegaly | description:"Cardiomegaly (PheCode 416)" | 19 | 45411941 | 0.6299 | 55106 |
| rs429358 | HypertrophicCardiomyopathy( | description:"Hypertrophic cardiomyopathy (sarcomere-positive)" | 19 | 45411941 | 0.655491 | 6699 |
| rs429358 | LVESI | description:"Left ventricular end systole inferior wall thickness" | 19 | 45411941 | 0.66 | 42094 |

| | | | | | | |
|---|---|---|---|---|---|---|
| rs429358 | LVESIL | description:"Left ventricular end systole inferolateral wall thickness" | 19 | 45411941 | 0.7 | 41947 |
| rs429358 | RVEDVtoLVEDV | description:"Right ventricular end-diastolic volume to left ventricular er | 19 | 45411941 | 0.71 | 41135 |
| rs429358 | LVGCS | description:"Left ventricular global circumferential strain" | 19 | 45411941 | 0.74 | 36033 |
| rs429358 | PAD_neversmk | description:"Peripheral artery disease in never smokers" | 19 | 45411941 | 0.75 | 9016 |
| rs429358 | PAStraincm | description:"Short-axis pulmonary artery strain (cm)" | 19 | 45411941 | 0.76 | 41135 |
| rs429358 | AVAP_AVRT | description:"Atrioventricular accessory pathways or atrioventricular rec | 19 | 45411941 | 0.794 | 11223 |
| rs429358 | LVEDAL | description:"Left ventricular end diastole anterolateral wall thickness" | 19 | 45411941 | 0.8 | 41913 |
| rs429358 | VaricoseVeins | description:"Varicose veins" | 19 | 45411941 | 0.8212 | 165034 |
| rs429358 | RVEF | description:"Right ventricular ejection fraction" | 19 | 45411941 | 0.83 | 41135 |
| rs429358 | AFlutter | description:"Atrial flutter (PheCode 427.22)" | 19 | 45411941 | 0.8339 | 106745 |
| rs429358 | PAH | description:"Pulmonary arterial hypertension" | 19 | 45411941 | 0.8472 | 6859 |
| rs429358 | LATEF | description:"Left atrial total emptying fraction" | 19 | 45411941 | 0.91 | 35648 |
| rs429358 | LVWTMax | description:"Left ventricular wall thickness (maximal)" | 19 | 45411941 | 0.91 | 36203 |
| rs429358 | RAMaxArea | description:"Right atrial maximum area" | 19 | 45411941 | 0.93 | 41135 |
| rs429358 | LVEDI | description:"Left ventricular end diastole inferior wall thickness" | 19 | 45411941 | 0.95 | 42122 |

**GWAS Sources**

| Dataset | Publication title | PMID |
|---|---|---|
| description:"Coronary artery disease (CAD)" | Discovery and systematic characteriz | PMID: 36474045 |
| description:"Abdominal aortic aneurysm" | Genome-wide association meta-anal | PMID: 37845353 |
| description:"Myocardial infarction" | Genome-wide analysis identifies nov | PMID: 33532862 |
| description:"Venous Thromboembolism" | Genomic Landscape of Thrombosis R | PMID: 39677447 |
| description:"Coronary artery plaque burden" | A genome-wide association study of | PMID: 40164586 |
| description:"Chronic pulmonary heart disease | Diversity and Scale: Genetic Architec | PMID: 37425708 |
| description:"Non-ischemic heart failure" | Genome-wide association study met | PMID: 40038546 |
| description:"Native myocardial T1 time" | Non-invasive Assessment of Organ-S | PMID: 38806679 |
| description:"Left ventricular end-systolic volur | Genetic analysis of right heart struct | PMID: 35697867 |
| description:"Left ventricular end-diastolic volu | Genetic analysis of right heart struct | PMID: 35697867 |

**Table S2.** Proteins associated with ApoER2

| E3>E4 | E4>E3 |
|---|---|
| AAK1 | A2M |
| ABCF1 | ABCE1 |
| ABCF2 | ACAA2 |
| ACADVL | ACAT1 |
| ACOT9 | ACLY |
| ACSL3 | ACO1 |
| ACSL4 | ACOT7 |
| ADAM9 | ACTB |
| AFAP1 | ACTC1 |
| AFG2A | ACTN1 |
| AGPS | ACTN4 |
| AHNAK | ACTR1A |
| ALYREF | ACTR2 |
| ANAPC7 | ACTR3 |
| AP1B1 | ADRM1 |
| AP2A1 | ADSL |
| AP2A2 | AFDN |
| AP2B1 | AGFG1 |
| AP2M1 | AGO2 |
| AP2S1 | AHCY |
| AP3B1 | AHSG |
| AP3D1 | AIMP1 |
| AP3M1 | AK2 |
| APEX1 | AKAP12 |
| ARAP3 | AKR1C3 |
| ARFGAP3 | ALB |
| ARHGAP29 | ALCAM |
| ARHGEF15 | ALDH18A1 |
| ARHGEF2 | ALDOA |
| ARL6IP1 | ANKHD1 |
| ARL6IP4 | ANP32A |
| ARL6IP5 | ANP32B |
| ARPC2 | ANXA1 |
| ARPC3 | ANXA2 |
| ARPC5 | ANXA5 |
| ARPC5L | ANXA6 |
| ASCC1 | ANXA7 |
| ASCC2 | APMAP |
| ASCC3 | APOE |
| ASPH | APOL2 |
| ATAD3A | ARCN1 |
| ATL3 | ARF1 |
| ATP1A1 | ARF4 |

| | |
|---|---|
| ATP5F1C | ARF6 |
| ATP5F1E | ARHGAP17 |
| ATP5MF | ARHGDIA |
| ATP5MG | ARHGDIB |
| ATP5PD | ARHGEF7 |
| ATP5PO | ARL8B |
| BCAP29 | ARPC1A |
| BCLAF1 | ARPC1B |
| BMS1 | ARPC4 |
| BMX | ATP1B3 |
| BUB3 | ATP2A2 |
| BYSL | ATP5F1A |
| BZW2 | ATP5F1B |
| C4A | ATP5PB |
| CALD1 | ATXN2 |
| CAVIN1 | ATXN2L |
| CAVIN2 | B2M |
| CAVIN3 | BASP1 |
| CCAR2 | BCAP31 |
| CCN1 | BCKDK |
| CCT7 | BTF3 |
| CD44 | BZW1 |
| CD93 | C11orf98 |
| CDC42BPB | CACYBP |
| CDC5L | CAD |
| CDC73 | CALM3 |
| CDK1 | CALR |
| CDK5RAP3 | CALU |
| CIRBP | CAMSAP2 |
| CKAP4 | CAND1 |
| CKAP5 | CANX |
| CLPB | CAP1 |
| CNBP | CAPN1 |
| COL18A1 | CAPN2 |
| CORO1C | CAPNS1 |
| COX4I1 | CAPRIN1 |
| COX5A | CAPZA1 |
| COX5B | CAPZA2 |
| CPNE3 | CAPZB |
| CPSF6 | CAST |
| CPSF7 | CAV1 |
| CPT1A | CBX3 |
| CRIP2 | CC2D1A |
| CS | CCDC124 |
| CSNK1A1 | CCN2 |
| CSNK1E | CCT2 |
| CSNK2B | CCT3 |

| | |
|---|---|
| CSRP1 | CCT4 |
| CTR9 | CCT5 |
| CYB5R3 | CCT6A |
| CYCS | CCT8 |
| CYP51A1 | CD151 |
| DBN1 | CD59 |
| DDOST | CD9 |
| DDX1 | CDC42 |
| DDX17 | CDC42BPA |
| DDX28 | CDK17 |
| DDX3X | CEP170 |
| DDX46 | CFAP20 |
| DDX5 | CFL1 |
| DDX6 | CFL2 |
| DECR2 | CLASP1 |
| DHCR24 | CLASP2 |
| DHCR7 | CLIC1 |
| DHX15 | CLIC4 |
| DHX36 | CLINT1 |
| DHX38 | CLTC |
| DHX57 | CNN2 |
| DHX9 | CNN3 |
| DIMT1 | CNOT1 |
| DNAJA1 | CNOT2 |
| DNAJA2 | CNP |
| DNAJA3 | COLGALT1 |
| DNAJB4 | COMT |
| DNAJC1 | COPA |
| DNAJC10 | COPB1 |
| DNAJC13 | COPB2 |
| DNAJC21 | COPE |
| DNAJC3 | COPG1 |
| DNM2 | COPS3 |
| DOCK10 | CSDE1 |
| DOCK4 | CSE1L |
| DOCK6 | CTNNA1 |
| DOCK7 | CTNNB1 |
| DPM1 | CTNND1 |
| DPM3 | CTSB |
| DSG1 | CTTN |
| DSP | CYFIP1 |
| DYNC1H1 | DARS1 |
| DYNLL1 | DECR1 |
| DYSF | DHX29 |
| ECE1 | DHX30 |
| ECH1 | DNAJB1 |
| ECI2 | DNAJC2 |

| | |
|---|---|
| EDF1 | DOCK1 |
| EEF1A1 | DOCK9 |
| EEF2 | DPYSL2 |
| EFTUD2 | DRG1 |
| EHD1 | DRG2 |
| EHD2 | DST |
| EHD4 | DSTN |
| EIF1 | DYNC1I2 |
| EIF1AX | DYNC1LI2 |
| EIF2A | EDC3 |
| EIF2D | EEF1B2 |
| EIF2S1 | EEF1D |
| EIF2S2 | EEF1E1 |
| EIF2S3 | EEF1G |
| EIF3A | EEFSEC |
| EIF3B | EIF2AK2 |
| EIF3CL | EIF2B1 |
| EIF3D | EIF2B2 |
| EIF3E | EIF2B3 |
| EIF3F | EIF2B4 |
| EIF3G | EIF2B5 |
| EIF3H | EIF4A1 |
| EIF3I | EIF4A2 |
| EIF3J | EIF4E2 |
| EIF3K | EIF4H |
| EIF3L | EIF5 |
| EIF3M | EIF5A |
| EIF4A3 | EIF6 |
| EIF4B | ELMO2 |
| EIF4E | ELOB |
| EIF4G1 | EMC2 |
| EIF4G2 | EML4 |
| EIF5B | ENG |
| EIPR1 | ENO1 |
| ELAVL1 | EPRS1 |
| ELMO1 | ERC1 |
| ELOA | ERLIN1 |
| EMC1 | ESM1 |
| EML3 | ESYT1 |
| EPHA2 | EXOC1 |
| EPHX1 | EXOC2 |
| ERGIC1 | EXOC3 |
| ERH | EXOC4 |
| ETF1 | EXOC5 |
| EWSR1 | EXOC6 |
| EXOSC10 | EXOC6B |
| EXOSC2 | EXOC7 |

| | |
|---|---|
| EZR | FABP5 |
| FADS2 | FAM107B |
| FAM98A | FAM120A |
| FANCI | FAM133B |
| FARSB | FAM91A1 |
| FAU | FAM98B |
| FBL | FAR1 |
| FDFT1 | FAR2 |
| FEN1 | FARSA |
| FERMT3 | FASN |
| FGD5 | FDPS |
| FHL1 | FDXR |
| FKBP11 | FH |
| FLG2 | FHL2 |
| FLII | FKBP3 |
| FMNL2 | FKBP4 |
| FMR1 | FLNA |
| FN1 | FLNB |
| FOCAD | FLNC |
| FRG1 | FMNL3 |
| FUS | FSCN1 |
| G3BP2 | FUBP1 |
| GCN1 | FUBP3 |
| GIMAP1 | FXR1 |
| GIMAP4 | FXR2 |
| GIMAP8 | G3BP1 |
| GIT2 | G6PD |
| GLG1 | GALNT1 |
| GMPR2 | GALNT2 |
| GNAQ | GANAB |
| GOLGA2 | GAPDH |
| GPX4 | GARS1 |
| GPX8 | GART |
| GRAMD1A | GBE1 |
| GRB10 | GCLM |
| GRN | GDI2 |
| GTF2E2 | GET3 |
| GTF2F1 | GFPT1 |
| GTPBP10 | GIGYF2 |
| H1-3 | GIT1 |
| H2AC11 | GLUD1 |
| H2AC21 | GMPS |
| H2BC11 | GNA11 |
| H2BC12 | GNAI2 |
| H3-3A | GNAS |
| H3-7 | GNB1 |
| H4C1 | GNB2 |

| | |
|---|---|
| HACD3 | GNG12 |
| HADHA | GNG5 |
| HCCS | GPI |
| HCFC1 | GPX1 |
| HDLBP | GSK3B |
| HERC1 | GSN |
| HHIP | GSTK1 |
| HMGA1 | GSTO1 |
| HMGB2 | GSTP1 |
| HMGB3 | GTF2F2 |
| HMOX1 | GTF2I |
| HNRNPA0 | GTPBP1 |
| HNRNPA1 | HADHB |
| HNRNPA2B | HARS1 |
| HNRNPA3 | HBA1 |
| HNRNPC | HBD |
| HNRNPD | HBS1L |
| HNRNPL | HEATR5B |
| HNRNPM | HLA-A |
| HNRNPR | HLA-B |
| HNRNPU | HMGB1 |
| HNRNPUL1 | HNRNPF |
| HNRNPUL2 | HNRNPH1 |
| HRNR | HNRNPH2 |
| HSD17B4 | HNRNPK |
| HSPA14 | HNRNPLL |
| HSPG2 | HSBP1 |
| HYAL2 | HSD17B12 |
| ICAM2 | HSP90AA1 |
| IDH2 | HSP90AB1 |
| IFI16 | HSP90B1 |
| IGF2BP2 | HSPA1B |
| IGF2BP3 | HSPA4 |
| IGHG3 | HSPA5 |
| IKBIP | HSPA8 |
| ILF2 | HSPA9 |
| ILF3 | HSPB1 |
| INF2 | HSPD1 |
| INPP5D | HSPE1 |
| IQGAP1 | HTRA1 |
| IQGAP3 | HYOU1 |
| KCTD12 | IARS1 |
| KHDRBS1 | IDH1 |
| KIF2A | IDH3A |
| KIF5B | IDH3B |
| KLC1 | IGFBP7 |
| KPLCE | ILK |

| | |
|---|---|
| KPNA2 | ILVBL |
| KTN1 | IMPDH2 |
| LARP1 | IPO5 |
| LARP4 | ITGA5 |
| LARS1 | ITGB1 |
| LBR | ITIH2 |
| LETM1 | JAK1 |
| LIMS1 | KARS1 |
| LMAN2 | KHSRP |
| LMF2 | KIF1B |
| LMNA | KIF1C |
| LPCAT2 | KPNB1 |
| LRP8 | LACTB |
| LRRC47 | LARP4B |
| LRRC59 | LASP1 |
| LRRC8A | LDHA |
| LSM6 | LDHB |
| LSM8 | LGALS1 |
| LTBP2 | LIMA1 |
| LTN1 | LIMS2 |
| LTV1 | LIMS4 |
| LUC7L2 | LMAN1 |
| LUZP1 | LOXL2 |
| MACF1 | LRRC8C |
| MAP1S | LRRFIP1 |
| MAP3K20 | LSG1 |
| MAP4K4 | LSM12 |
| MAP4K5 | LSS |
| MAP7D1 | LUC7L |
| MARK2 | LUC7L3 |
| MATR3 | MANF |
| MBNL1 | MAP1B |
| MBOAT7 | MAP2 |
| MCM3 | MAP4 |
| MCM4 | MAPK1 |
| MCM5 | MAPRE1 |
| METAP2 | MARCKS |
| MFAP1 | MARK3 |
| MLST8 | MARS1 |
| MOGS | MAT2B |
| MOV10 | MCAM |
| MRM3 | MCTS1 |
| MRPL1 | MDH2 |
| MRPL10 | MINK1 |
| MRPL11 | MLEC |
| MRPL13 | MLKL |
| MRPL14 | MMP1 |

| | |
|---|---|
| MRPL15 | MMP14 |
| MRPL16 | MMRN1 |
| MRPL17 | MMRN2 |
| MRPL18 | MMTAG2 |
| MRPL19 | MRPL37 |
| MRPL20 | MSN |
| MRPL21 | MT2A |
| MRPL22 | MTHFD1L |
| MRPL27 | MTOR |
| MRPL28 | MTREX |
| MRPL3 | MTSS1 |
| MRPL30 | MVP |
| MRPL38 | MYDGF |
| MRPL39 | MYH9 |
| MRPL43 | MYL12B |
| MRPL44 | MYL6 |
| MRPL47 | MYO1E |
| MRPL49 | MYO5A |
| MRPL53 | MYO9B |
| MRPL9 | MZT2B |
| MRPS18A | NAA10 |
| MRPS22 | NAA15 |
| MRPS9 | NAA50 |
| MRTO4 | NACA |
| MSH6 | NANS |
| MT-CO2 | NAP1L1 |
| MTDH | NAP1L4 |
| MTHFD1 | NARS1 |
| MTMR10 | NASP |
| MYCBP2 | NAV1 |
| MYCT1 | NEDD1 |
| MYO1C | NEMF |
| MYO1D | NES |
| MYO6 | NIBAN2 |
| MYOF | NME2 |
| NAMPT | NNMT |
| NCAPD2 | NSF |
| NCKAP1 | NSUN2 |
| NCL | NUCKS1 |
| NDUFA4 | NUDC |
| NEK7 | NUFIP2 |
| NELFB | OLA1 |
| NHERF2 | OXSR1 |
| NIPSNAP2 | P3H1 |
| NMT1 | P4HB |
| NMT2 | PABPC1 |
| NOB1 | PABPC4 |

| | |
|---|---|
| NOMO1 | PAFAH1B1 |
| NONO | PAICS |
| NOP56 | PARK7 |
| NOP58 | PARVA |
| NOSIP | PAWR |
| NPM1 | PCBP1 |
| NPM3 | PCBP2 |
| NQO1 | PCNA |
| NSDHL | PCNP |
| NUDT21 | PDCD6IP |
| OAS3 | PDIA3 |
| OCIAD2 | PDIA4 |
| OSBPL3 | PDIA6 |
| PA2G4 | PDLIM1 |
| PAF1 | PEBP1 |
| PAK4 | PFKL |
| PAPSS1 | PFN1 |
| PAPSS2 | PGAM1 |
| PARP4 | PGD |
| PBXIP1 | PGK1 |
| PDAP1 | PICALM |
| PDCD4 | PIGK |
| PDIA5 | PIR |
| PDLIM5 | PKM |
| PDLIM7 | PLCB3 |
| PECAM1 | PLG |
| PFKM | PLIN3 |
| PFKP | PLOD2 |
| PGAM5 | PLS3 |
| PHB1 | PNP |
| PHB2 | PNPT1 |
| PHF6 | PPA1 |
| PHLDB1 | PPIA |
| PIK3C2A | PPIB |
| PIK3CB | PPM1G |
| PKN2 | **PPP2CA** |
| PLEC | **PPP2R1A** |
| PLOD1 | **PPP2R2A** |
| PLOD3 | PRDX1 |
| PNO1 | PRDX2 |
| PNPLA6 | PRDX3 |
| POLR2A | PRDX6 |
| POLR2B | PRKCSH |
| POLR2C | PRKRA |
| POLR2E | PRPS1 |
| POSTN | PRRC2A |
| PPFIBP1 | PRRC2C |

| | |
|---|---|
| PPIL4 | PSMC1 |
| PPP1CA | PSMC4 |
| PPP1CB | PSMD13 |
| PPP1R12A | PSMD2 |
| PREX1 | PSMD3 |
| PRKCH | PSMD7 |
| PRKDC | PSMD8 |
| PRMT5 | PSME1 |
| PRPF19 | PTBP1 |
| PRPF3 | PTGES3 |
| PRPF31 | PTMA |
| PRPF4 | PTPN14 |
| PRPF6 | PTPRB |
| PRPF8 | PUM1 |
| PTBP3 | PVR |
| PTK2 | PXDN |
| PTPN1 | PXN |
| PUF60 | PYCR2 |
| PYCR1 | QARS1 |
| PYM1 | RAB11B |
| RAB12 | RAB11FIP5 |
| RAB35 | RAB13 |
| RAB8A | RAB14 |
| RAB8B | RAB1A |
| RABL6 | RAB1B |
| RACK1 | RAB5C |
| RAI14 | RAC1 |
| RALB | RAC2 |
| RALY | RALA |
| RBM17 | RALGAPA1 |
| RBM3 | RAN |
| RBM39 | RANBP1 |
| RBMX | RANGAP1 |
| RDH11 | RAP1B |
| RDX | RAPH1 |
| RECQL | RARS1 |
| REEP5 | RASIP1 |
| RIOK1 | RBBP7 |
| RIOK2 | RBM25 |
| RNF213 | RBMS2 |
| ROCK2 | RCC2 |
| RPL10 | RCN1 |
| RPL13A | RHOA |
| RPL14 | RHOG |
| RPL15 | RNH1 |
| RPL17 | RPA1 |
| RPL18 | RPL10A |

| | |
|---|---|
| RPL18A | RPL11 |
| RPL19 | RPL12 |
| RPL21 | RPL13 |
| RPL22 | RPL22L1 |
| RPL23A | RPL23 |
| RPL24 | RPL27 |
| RPL26 | RPL27A |
| RPL26L1 | RPL29 |
| RPL28 | RPL31 |
| RPL3 | RPL35 |
| RPL30 | RPL37 |
| RPL32 | RPL5 |
| RPL34 | RPL8 |
| RPL35A | RPLP2 |
| RPL36 | RRAS |
| RPL36A | RRM1 |
| RPL36AL | RSU1 |
| RPL37A | RTN4 |
| RPL38 | RUVBL1 |
| RPL39 | RUVBL2 |
| RPL4 | S100A11 |
| RPL6 | S100A13 |
| RPL7 | S100A6 |
| RPL7A | SAE1 |
| RPL9 | SAR1A |
| RPLP0 | SCCPDH |
| RPN1 | SCP2 |
| RPN2 | SDCBP |
| RPS10 | SEC23A |
| RPS11 | SEC31A |
| RPS12 | SEPTIN8 |
| RPS13 | SERPINE1 |
| RPS14 | SETSIP |
| RPS15 | SH3BGRL3 |
| RPS15A | SHANK3 |
| RPS16 | SHMT2 |
| RPS17 | SKIC2 |
| RPS18 | SLAIN2 |
| RPS19 | SLC16A1 |
| RPS2 | SMAD3 |
| RPS20 | SMC2 |
| RPS21 | SMG9 |
| RPS23 | SMURF2 |
| RPS24 | SQSTM1 |
| RPS25 | SRP54 |
| RPS26 | SRP68 |
| RPS27 | SRP72 |

| | |
|---|---|
| RPS27A | SRPK2 |
| RPS27L | SRSF11 |
| RPS28 | SSB |
| RPS29 | ST13 |
| RPS3 | STAU1 |
| RPS3A | STIP1 |
| RPS4X | STMN1 |
| RPS5 | STRAP |
| RPS6 | STX12 |
| RPS7 | STX7 |
| RPS8 | SUB1 |
| RPS9 | SWAP70 |
| RPSA | SYNJ2 |
| RRBP1 | TAF15 |
| RRP12 | TAGLN2 |
| RSRC2 | TALDO1 |
| RTCB | TAOK1 |
| RTRAF | TAOK2 |
| S100A10 | TARS1 |
| S100A16 | TBC1D4 |
| SACM1L | TCP1 |
| SACS | TGM2 |
| SART1 | TIA1 |
| SART3 | TJP2 |
| SBDS | TKT |
| SDAD1 | TLN1 |
| SDHA | TMED5 |
| SEC22B | TMED9 |
| SEC61A1 | TMEM214 |
| SEC61B | TMSB10 |
| SEC63 | TMSB4X |
| SEPTIN11 | TMX1 |
| SEPTIN2 | TNKS1BP1 |
| SEPTIN7 | TNPO1 |
| SERBP1 | TPD52L2 |
| SERF2 | TPI1 |
| SERPINH1 | TPM3 |
| SF3B1 | TPM4 |
| SF3B2 | TPP1 |
| SFPQ | TPT1 |
| SFXN1 | TRAP1 |
| SH3BP4 | TRAPPC5 |
| SH3PXD2B | TRIM22 |
| SKIC3 | TRIM28 |
| SKIC8 | TRIP4 |
| SLC25A1 | TUBA1A |
| SLC25A3 | TUBA1B |

| | |
|---|---|
| SLC25A5 | TUBB |
| SLC25A6 | TUBB2A |
| SMC3 | TUBB3 |
| SMC4 | TUBB4B |
| SMG1 | TUBB6 |
| SMG8 | TUBG1 |
| SND1 | TUBGCP2 |
| SNRNP200 | TUBGCP3 |
| SNRNP40 | TUFM |
| SNRNP70 | TXN |
| SNRPB | TXNDC5 |
| SNRPD1 | TXNRD1 |
| SNRPD2 | U2AF1 |
| SNRPD3 | UBA1 |
| SNU13 | UBAP2 |
| SNX9 | UBE2D3 |
| SPATS2 | UBE2N |
| SPATS2L | UBR4 |
| SPCS2 | UCHL1 |
| SPCS3 | UGDH |
| SPTLC1 | UNC45A |
| SQOR | UQCRC1 |
| SRGAP2 | UQCRC2 |
| SRP14 | VAC14 |
| SRP19 | VAPB |
| SRP9 | VAT1 |
| SRPK1 | VCL |
| SRPRA | VCP |
| SRPRB | VIM |
| SRRM1 | VPS11 |
| SRSF1 | VWF |
| SRSF2 | WARS1 |
| SRSF3 | WBP11 |
| SRSF4 | WDR1 |
| SRSF5 | XRCC5 |
| SRSF6 | XRCC6 |
| SRSF7 | YARS1 |
| SSR4 | YARS2 |
| SSRP1 | YBX1 |
| STAB1 | YBX3 |
| STAU2 | YTHDC2 |
| STING1 | YWHAB |
| STK10 | YWHAE |
| STT3A | YWHAG |
| STT3B | YWHAH |
| STX4 | YWHAQ |
| SUCLG2 | YWHAZ |

SUPT16H
SURF4
SYNCRIP
SYNE2
SYNPO
TANC1
TAOK3
TARDBP
TCEA1
TCF25
TECR
THBS1
THRAP3
TJP1
TM9SF2
TMA7B
TMCO1
TMED10
TMED2
TMED7
TMEM109
TMEM33
TNIK
TOMM34
TOR1AIP1
TPP2
TRA2A
TRA2B
TRIM21
TRIM25
TRIM56
TRIOBP
TRMT10C
TSR1
TXLNA
U2AF2
UBAP2L
UBE2I
UBL5
UBXN4
UFL1
UPF1
UPF3B
USP10
USP16
USP24
USP39
ZNF598
ZRANB2
ZYX

USP7
USP8
VAPA
VASP
VDAC2
VPS53
VTI1B
WASL
WDR11
WDR77
WDR82
XRN1
XRN2
YTHDF2
YTHDF3
ZC2HC1A
ZC3H15
ZC3H4
ZC3H7B
ZC3HAV1
ZNF622

**Supplemental Videos**

**Video S1.** Related to Figure 2. IVM recording of thrombosis in the mesenteric microcirculation in an hE3 (humanized ApoE3) mouse.

**Video S2.** Related to Figure 2. IVM recording of thrombosis in the mesenteric microcirculation in an hE4 (humanized ApoE4) mouse.
